# Chromatin accessibility and histone acetylation in the regulation of competence in early development

**DOI:** 10.1101/797183

**Authors:** Melody Esmaeili, Shelby A. Blythe, John W. Tobias, Kai Zhang, Jing Yang, Peter S. Klein

**Affiliations:** Cell and Molecular Biology Graduate Group, University of Pennsylvania Perelman School of Medicine, Philadelphia, PA, USA; Department of Molecular Biosciences, Northwestern University, Evanston, IL, USA; Penn Genomic Analysis Core and Abramson Cancer Center, University of Pennsylvania, Perelman School of Medicine, Philadelphia, PA, USA; Department of Biochemistry, University of Illinois at Urbana-Champaign, Urbana, IL, USA; Department of Comparative Biosciences, University of Illinois at Urbana-Champaign, Urbana, IL, USA; Departments of Medicine (Hematology-Oncology) and Cell and Developmental Biology, University of Pennsylvania Perelman School of Medicine, Philadelphia, PA, USA

**Keywords:** competence, induction, Xenopus, dorsal development, ATAC-Seq, chromatin

## Abstract

As development proceeds, inductive cues are interpreted by competent tissues in a spatially and temporally restricted manner. While key inductive signaling pathways within competent cells are well-described at a molecular level, the mechanisms by which tissues lose responsiveness to inductive signals are not well understood. Localized activation of Wnt signaling before zygotic gene activation in *Xenopus laevis* leads to dorsal development, but competence to induce dorsal genes in response to Wnts is lost by the late blastula stage. We hypothesize that loss of competence is mediated by changes in histone modifications leading to a loss of chromatin accessibility at the promoters of Wnt target genes. We use ATAC-seq to evaluate genome-wide changes in chromatin accessibility across several developmental stages. Based on overlap with p300 binding, we identify thousands of putative cis-regulatory elements at the gastrula stage, including sites that lose accessibility by the end of gastrulation and are enriched for pluripotency factor binding motifs. Dorsal Wnt target gene promoters are not accessible after the loss of competence in the early gastrula while genes involved in mesoderm and neural crest development maintain accessibility at their promoters. Inhibition of histone deacetylases increases acetylation at the promoters of dorsal Wnt target genes and extends competence for dorsal gene induction by Wnt signaling. Histone deacetylase inhibition, however, is not sufficient to extend competence for mesoderm or neural crest induction. These data suggest that chromatin state regulates the loss of competence to inductive signals in a context-dependent manner.

## Introduction

The ability of developing tissues to interpret and effect change in response to inductive cues is termed competence (Waddington, 1940). Using heterochronic transplants of the amphibian dorsal organizer into host embryos at different stages of development, Spemann, Mangold, and colleagues demonstrated the fundamental principle that embryonic tissues respond to inductive signals within temporally and spatially restricted windows of competence (Hamburger, 1988; Nakamura et al., 1978; Spemann, 1938). This control of competence is essential for cell fate specification and patterning and enables embryos to reuse a limited number of signaling pathways for distinct outcomes throughout development. The molecular mechanisms of signaling by pathways such as Wnt, TGF-ß, FGF, and others have been worked out in detail yet the mechanisms by which tissues gain and lose competence to respond to these signals are incompletely understood.

One of the earliest inductive processes in amphibian development is the specification of the dorsal-ventral axis through localized activation of Wnt signaling (De Robertis et al., 2000; Moon and Kimelman, 1998; Sokol, 1999). In *Xenopus laevis*, as well as zebrafish, Wnt signaling is initiated before zygotic genome activation through the activity of maternal factors, including localized Wnts (Cha et al., 2008; Lu et al., 2011; Tao et al., 2005) and *Huluwa* (Yan et al., 2018), resulting in localized transcription of dorsal-specifying genes after the midblastula transition (MBT). The paired box transcription factor *siamois* (*sia1 and sia2*), a major direct target of early Wnt signaling (Laurent et al., 1997; Lemaire et al., 1995), is both necessary and sufficient for subsequent dorsal development (Bae et al., 2011; Ding et al., 2017; Ishibashi et al., 2008; Kessler, 1997). While the entire equatorial region (marginal zone) of the embryo is competent to respond to Wnt signaling during early cleavage stages, equatorial cells that do not receive a Wnt signal during cleavage stages contribute to future ventral and lateral mesoderm. These future ventral and lateral cells can be induced to express dorsal genes, such as *sia1* and *nodal3.1* (previously known as *Xnr3)*, by ectopic activation of the Wnt pathway (Ding et al., 2017), but competence for this response declines after the 64-cell stage and is lost after the MBT (Blythe et al., 2010; Christian and Moon, 1993; Darken and Wilson, 2001; Hamilton et al., 2001; Kao et al., 1986; Yamaguchi and Shinagawa, 1989; Yang et al., 2002). Although future ventral and lateral cells progressively lose the ability to express dorsal genes in response to Wnt signaling, the daughters of these cells still respond to Wnts during later stages, for example in response to anterior-posterior patterning signals (Christian and Moon, 1993; Fredieu et al., 1997; Hamilton et al., 2001; Kjolby and Harland, 2017; Nakamura et al., 2016; Yamaguchi and Shinagawa, 1989).

The differential response to Wnt signaling during early stages of *Xenopus* development has been examined in greater depth recently. A genome-wide approach identified 123 genes that are enriched on the dorsal side of gastrula stage embryos and are activated by the maternal Wnt/ß-catenin signaling pathway (Ding et al., 2017). Remarkably, only 3 of these overlapped with the set of Wnt-dependent genes activated by zygotic/early gastrula Wnt signaling in *X. laevis* (Kjolby and Harland, 2017) or *X. tropicalis* (Nakamura et al., 2016). That the same signaling pathway yields distinct developmental outcomes within a given population of cells at times separated by only a few hours suggests that the signal transduction mechanisms remain intact and that the loss of competence likely occurs at the level of target gene transcription (Darken and Wilson, 2001; Hamilton et al., 2001).

The precise temporal and spatial control of gene expression during development necessitates precise regulation of chromatin organization including changes in chromatin accessibility, histone modifications, and DNA methylation. Generally, transcription occurs in regions of the genome that are physically accessible (“open”) to transcription factors (Shahbazian and Grunstein, 2007). Conversely, inactive genes are typically associated with inaccessible (“closed”) chromatin (Beisel and Paro, 2011). Recent work in diverse organisms from *Drosophila* to vertebrates has identified transcription factors involved in the earliest steps in the activation of zygotic genes. In *Xenopus*, for example, Wnt signaling directs the arginine methyltransferase PRMT2 to dorsal Wnt target genes prior to zygotic gene activation (ZGA), establishing a poised chromatin state at promoters that become active at later stages, once global zygotic gene expression has been activated (Blythe et al., 2010). Later in the blastula stage, maternal transcription factors including Fox, Sox, and Oct/Pou family members modify chromatin more globally to establish competence for primary germ layer formation (Charney et al., 2017; Gentsch et al., 2019; Paraiso et al., 2019). In a role analogous to maternal transcription factors Zelda and Grainyhead in *Drosophila* (Jacobs et al., 2018; McDaniel et al., 2019), maternal Sox and Oct/Pou family transcription factors are required for activation of zygotic gene expression in zebrafish and *X. tropicalis* (Gentsch et al., 2019; Lee et al., 2013; Leichsenring et al., 2013; Paraiso et al., 2019), functioning as pioneer factors that confer competence for inductive signals at the time of zygotic gene activation (Gentsch et al., 2019).

However, the mechanisms by which changes in chromatin architecture mediate loss of competence to early inductive signals, including dorsal Wnt signaling, remain unclear. We hypothesize that loss of competence is associated with a loss of chromatin accessibility. We used Assay for Transposase-Accessible Chromatin followed by sequencing (ATAC-Seq) (Buenrostro et al., 2013) to examine chromatin accessibility at stages when competence to respond to several inductive signals is changing, including induction of the dorsal organizer, mesoderm, and neural crest. Our data support the hypothesis that loss of competence to Wnt signaling to induce dorsal development is correlated with chromatin inaccessibility and is mediated by histone deacetylation at dorsal Wnt target gene promoters.

## Materials and Methods

### Experimental model and vertebrate animal handling

In vitro fertilization, microinjection and culture of *Xenopus laevis* embryos were performed as described (Sive et al., 2000). *Xenopus* embryos were obtained by in vitro fertilization and cultured at room temperature in 0.1X MMR buffer. Embryo staging was based on Nieuwkoop and Faber (Nieuwkoop and Faber, 1967) as reproduced in Xenbase (http://www.xenbase.org/, RRID:SCR_003280), which also served as a reference for gene expression patterns, gene sequences, and sequence alignments. *Xenopus* experiments were approved by the Institutional Animal Care and Use Committee at the University of Pennsylvania.

### Embryo culture

For dorsal induction, embryos were treated at the 32-64 cell stage with 0.3M LiCl for 10-12 minutes, followed by 3 washes in 0.1xMMR. A separate group of embryos at the 32-64 cell stage was placed into 0.1XMMR with 100nM TSA and cultured until late blastula stage (Nieuwkoop and Faber stage 9) and then half of the TSA-treated group and an equal number of untreated control embryos were exposed to 0.3M LiCl for 12-14 minutes followed by 3 washes in 0.1xMMR. The TSA treated embryos were returned to 0.1XMMR with 100nM TSA. At the onset of gastrulation, embryos were washed 3 times in 0.1xMMR and frozen at −80°C in minimal buffer. For these whole embryo experiments and explant experiments described below, sibling embryos were cultured until the tadpole stage to assess morphological effects of LiCl exposure; embryos or explants were only processed for further analysis if embryos exposed to LiCl in cleavage stage were dorsalized (defined as ≥ 80% of sibling tadpoles showing complete lack of trunk structures, no obvious somites, expanded eyes, and frequently expanded cement gland (dorsal-anterior index (DAI) ≥ 8; (Kao and Elinson, 1988; Karimi et al., 2018)) and embryos exposed to LiCl at the late blastula stage demonstrated anterior truncation/posteriorization (defined as ≥ 80% of sibling tadpoles showing reduced or absent forebrain and small or absent cement gland (DAI ≤ 4), frequently with small or absent eyes and microcephaly.

Explant assays: For ventral marginal zone explants (VMZ), embryos were treated with LiCl and/or TSA as above, rinsed in 0.1xMMR at the onset of gastrulation, and transferred to 0.1XMMR at 12°C. VMZs were dissected using a Gastromaster and then immediately frozen. To measure induction of *sia1* and *nodal3.1* in ectodermal explants (animal caps), embryos were treated as above and transferred to 12°C at the onset of gastrulation. Animal cap explants were dissected in 0.5xMMR and immediately frozen for later RNA extraction. For mesoderm induction assays, animal caps were dissected at the late blastula or early gastrula stage and transferred to 0.5X MMR with 0.1%BSA and with or without bFGF (25ng/mL). A subset of embryos were treated with 100nM TSA from the late blastula stage until the onset of gastrulation and then animal caps were dissected and transferred to medium with or without FGF. Explants were cultured until sibling embryos reached the neurula stage (stage 13) and were then harvested for analysis of mesodermal gene expression. For neural crest induction assays, embryos were placed in 0.5X MMR, 3% Ficoll at 12°C and microinjected into each blastomere at the animal pole of the two-cell stage with mRNA encoding *chordin* (*chrd.1*; 0.5ng/blastomere) with or without mRNA encoding the HMG domain of Tcf3 fused to the VP16 activation domain and the glucocorticoid receptor ligand binding domain, termed THVGR (10pg/blastomere) (Wu et al., 2005; Yang et al., 2002). Embryos were allowed to develop until stage 9 when animal cap explants were dissected in 0.5XMMR using a Gastromaster. Explants were treated with control medium or 10mM dexamethasone in 0.5XMMR at stage 10. Separate groups of explants were placed into 100nM TSA or control medium at stage 10 and allowed to develop until stage 12.5 when they were treated with dexamethasone. At stage 14 explants were transferred to 12°C and cultured until siblings reached stage 22. Explants were rinsed three times in 0.5XMMR and frozen. For synthesis of *chrd* (Piccolo et al., 1996) and THVGR (Yang et al., 2002) mRNAs, templates were linearized from pCS2-based plasmids and capped mRNA was synthesized using the mMessage mMachine SP6 Transcription Kit (ThermoFisher AM1340) and purified using an RNeasy Mini Kit (QIAGEN).

### RNA isolation, cDNA synthesis, and reverse transcription-quantitative PCR (RT-qPCR)

RNA was isolated from frozen embryos or explants using Qiagen RNeasy with on-column DNase (Qiagen On-column RNAse-free DNAse Set). Reverse transcription was performed using 1ug (whole embryos) or 0.1ug (explants) of RNA (Yang et al., 2002). Real-time PCR was performed with Power SYBR Green PCR Master Mix (ThermoFisher) using primers indicated in Table I. Ornithine decarboxylase (*ODC*) served as a reference gene. Data were analyzed using the Comparative CT Method (ΔΔCt) (Taneyhill and Adams, 2008): data were first normalized to *ODC* and then to an indicated control group to calculate fold increase. For ectodermal explant (animal cap) assays, expression of *sia1* in untreated control explants was frequently undetectable and we were not able to represent the induction of *sia1* as fold-increase relative to control explants for these samples; nevertheless, we observed robust induction of *sia1* (and *nodal3.1*) by LiCl in stage 9 embryos pre-treated with TSA as well as in control embryos exposed to LiCl at stage 6 in 9 out of 10 independent experiments. Figure 4B shows those experiments for which we could measure basal expression of *sia1* and report a fold-increase in the experimental groups; regression methods were used to assess all 10 independent explant experiments simultaneously (see “statistical analysis” below), which confirmed a significant increase in both *sia1* and *nodal3.1* expression when explants from TSA-treated embryos were exposed to lithium at stage 9, compared to lithium alone or TSA alone.

**Table 1.**
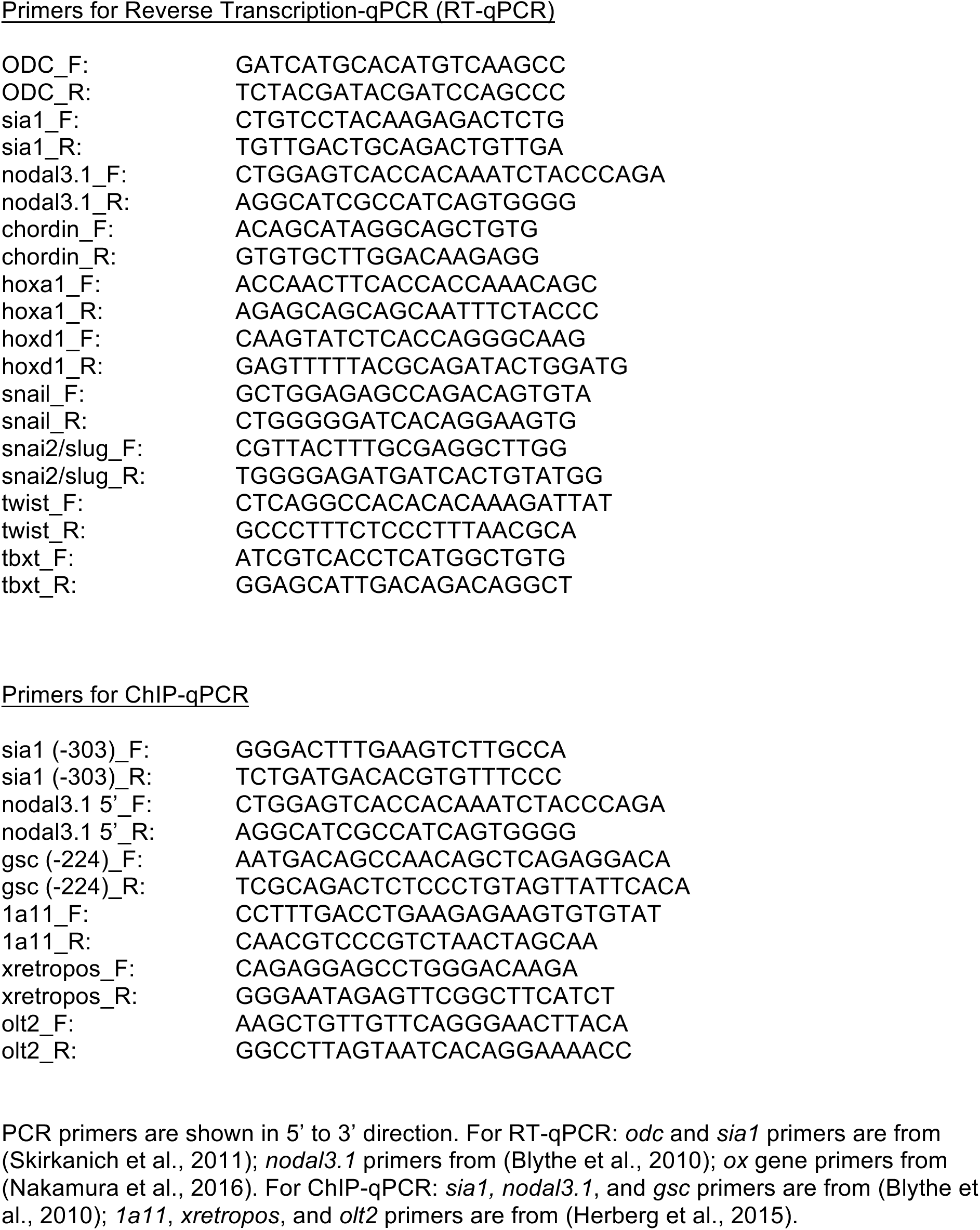
Primer Sequences

### Chromatin Immunoprecipitation

Chromatin immunoprecipitation was performed as in (Blythe et al., 2010; Blythe et al., 2009) with a few modifications. Embryos were fixed at stage 10 for 60 minutes when a Branson 250 sonifier was used to fragment chromatin or for 15 minutes when a Covaris sonicator was used. Embryos were treated with glycine, washed in PBS (Blythe et al., 2010; Blythe et al., 2009), and either frozen in minimal buffer or dissected into dorsal-ventral halves with a microblade attached to a surgical clamp using the dorsal blastopore lip as a landmark. Embryos or embryo halves were then frozen in minimal buffer and stored at −80°C. Elution of immunoprecipitated DNA from Protein A-agarose beads was followed by RNAseA for 1hr and proteinase K treatment with reversal of crosslinking overnight. DNA was resuspended in 100uL H20 and 1uL of DNA was used for real time-qPCR (ChIP-qPCR). Antibodies used for ChIP included: IgG (Abcam Ab171870); H3K9ac (Cell Signaling 9671S); H3K9me3 (Abcam Ab8898); H3K27me3 (EMD Millipore 07-449).

### ATAC-Seq

ATAC-Seq methods were adapted from (Buenrostro et al., 2013). *Xenopus laevis* ectodermal explants (animal caps) were dissected at the blastula stage and collected at the onset of gastrulation (stage 10) or at the end of gastrulation (stage 12). We empirically determined the optimal number of explants by analyzing nucleosome ladders by 6% PAGE (Buenrostro et al., 2013). The optimal sample size was 2 explants per 50 µl Tn5 reaction; we were unable to achieve adequate transposition reactions with explants prior to stage 9 or with whole embryos. Therefore, 2 explants per biological replicate (3 biological replicates per stage) were transferred to 1 ml of ice ice-cold PBS and centrifuged at 500xg for 5 minutes at 4°C. The supernatant was removed, 1 ml of PBS was added, explants were centrifuged again, and the supernatant was aspirated completely. Explants were resuspended in 50 µl RSB (10mM Tris pH 7.4, 10mM NaCl, 3mM MgCl2) with 0.1% (v/v) Igepal CA-630 at 4°C and dispersed by pipetting. The samples were centrifuged at 500xg for 10 minutes at 4°C. Supernatants were removed and pellets were resuspended in 1X TD buffer (10mM Tris pH 7.6, 5 mM MgCl2, and 10% dimethylformamide) and 2.5µL of Tn5 (3µM) was added. Samples were incubated at 37°C for 1 hour with constant shaking and digested with protease K overnight. DNA was purified with 1.8 volumes of Ampure XP Beads (Beckman Coulter), eluted in 40 µl, amplified by PCR with barcoded primers as described, purified using AMPure XP beads (Thermo-Fisher) to remove adaptors, and submitted in triplicate for sequencing on a HiSeq 4000 with 100 nucleotide paired-end reads.

### Data Analysis

Raw sequences were submitted to the ATAC-seq pipeline currently in use by the ENCODE project for mapping to the xenLae2 build of the *Xenopus laevis* genome (http://genome.ucsc.edu/cgi-bin/hgGateway?db=xenLae2) and peak detection (https://www.encodeproject.org/atac-seq/, version 1.4.2 of the code at https://github.com/ENCODE-DCC/atac-seq-pipeline). Samples were divided by stage, and each stage was submitted as a replicate set in two parallel runs using default parameters, including irreproducible discovery rate filtering (IDR threshold 0.05), with the exception of allowing for the retention of non-standard chromosome names and the allocation of high memory to accomodate the structure of the genome build.

Resulting peak coordinates for each replicate set were merged, and de-duplicated reads for each sample aligning to each region were counted with featureCounts (v1.6.5, http://subread.sourceforge.net)(Liao et al., 2014). The resulting table was analyzed for differential peak counts between the two stages with DESeq2 (v1.25.9, https://bioconductor.org/packages/release/bioc/html/DESeq2.html) (Love et al., 2014). Peaks were annotated for their relationship to known XenLae2 genomic features with HOMER (v4.10, http://homer.ucsd.edu/homer/)(Heinz et al., 2010) (Supplementary Data1). ATAC-Seq data were deposited in the Gene Expression Omnibus database (GSE138905). Functional annotation/gene ontology analyses were performed using DAVID 6.8 (Huang da et al., 2009); enriched Uniprot keywords with false discovery rate (FDR) ≤ 0.01 are shown in the figures.

Gene tracks were viewed using Integrative Genomics Viewer (IGV) software (Robinson et al., 2011). For comparison of ATAC reads and expression in stage 10 ectodermal explants, the set of genes that were annotated in both the Agilent microarray used by (Livigni et al., 2013) and in the xenLae2 genome build were used (Supplementary Data 2). Log_2_ of normalized mean ATAC reads (for values > 0) was plotted vs. Log_2_ normalized expression detected in the microarray. To compare ATAC-Seq to ChIP for p300, raw sequence for p300 ChIP (GSE76059) for duplicate samples from stage 10.5 embryos, was was subjected to peak-calling against the xenLae2 genome using HOMER, and ATAC-Seq peaks were annotated for their overlap with p300 peaks. ATAC-Seq tracks for selected genes in gastrula stage *Xenopus laevis* were compared to Dnase-Seq data from *Xenopus tropicalis* embryos at the midblastula stage (GSE113186) (Gentsch et al., 2019) using IGV software.

For motif analysis we defined the following peak lists: 1. pCRMs (putative cis-regulatory modules) were defined as accessible peaks (based on ATAC) associated with p300 (with ≥ 30% overlap) at stage 10 and that were annotated as intergenic or intronic (n=17,519 peaks), and 2. ΔpCRMs were defined as the subset of pCRMs associated with accessible peaks at stage 10 that were reduced at stage 12 such that Log_2_[fold change in mean counts at st10/st12] ≤ −1 with an adjusted p-value < 0.05 and which were within 100kb of a TSS (n=1897 peaks). Motif enrichment analysis was performed with HOMER (Heinz et al., 2010) using “-size given” and all other parameters as default. Three enrichment analyses were run, pCRM vs genome (background), ΔpCRM vs genome, and ΔpCRM vs a background of pCRM sequences.

### Statistical Analysis

All experiments were repeated a minimum of three times. Error bars in all cases represent standard error (SEM). In some cases, the magnitude of induction was variable and the standard deviation exceeded the mean values; however, in these cases, the experiments were qualitatively reproducible. To compare samples from the 10 biological replicates for the dorsal induction in ectodermal explant assays, we used generalized linear models fitted with generalized estimating equations (GEE) to compare ΔCt values in a 2×2 factorial design. We estimated main effects for lithium alone at stage 9, TSA treatment alone, and tested the interaction term for TSA+lithium. Significance was tested using the z-score corresponding to the model coefficient of interest. Means and p-values for the comparisons are reported in the legend to Figure 4B. The GEE method accounts for random effects as correlated outcomes within biological samples that are the same across treatments.

## Results

### Chromatin Accessibility at the onset of Gastrulation

We hypothesized that loss of competence to respond to inductive signals would be associated with inaccessibility of regulatory regions within stage-specific genes. We therefore assessed genome-wide chromatin accessibility using ATAC-Seq in ectodermal explants at the onset of gastrulation (Nieuwkoop and Faber stage 10). Although ATAC-Seq has been widely used in other organisms, and a method for ATAC-Seq in *X. tropicalis* was described recently (Bright and Veenstra, 2019), genome-wide ATAC-Seq data have not yet been reported for *Xenopus laevis* or *X. tropicalis* embryos for any stage of development. Given this, it was critical to optimize ATAC protocols for early Xenopus embryos; we were unable to obtain optimal quality chromatin digestion using whole embryos and suspected that a component of early embryos, likely yolk protein, interfered with transposase activity, as also reported by others (Gentsch et al., 2019). We found that optimal transposase digestion at early developmental stages occurred only with ectodermal explants, which have lower amounts of yolk per cell compared to whole embryos. In addition, it was essential to reduce the number of cells per reaction considerably below that typically used in standard ATAC protocols (see Methods). We chose to examine chromatin accessibility in ectodermal explants at the onset of gastrulation (stage 10), when competence to respond to dorsal and mesodermal inducing signals has been lost while competence for neural and neural crest induction persists (Dale et al., 1985; Darken and Wilson, 2001; Gurdon et al., 1985; Jones and Woodland, 1987; Kengaku and Okamoto, 1993; Kodjabachian and Lemaire, 2001; Lamb et al., 1993; Mancilla and Mayor, 1996).

We obtained 46 million paired reads mapping uniquely to the nuclear genome. ATAC-Seq on early gastrula stage (stage 10) ectodermal cells identified ∼70,000 peaks with 5720 (8%) peaks at annotated transcription start sites (TSS), 11,416 (16%) in intragenic regions, 278 at transcription termination sites, and the remainder distributed in intergenic or unannotated regions (Figure 1A, Supplementary Data 1). *Xenopus laevis* underwent a genome duplication approximately 17 million years ago; the resulting allotetraploid genome comprises two homeologous subgenomes termed L (long chromosomes) and S (short chromosomes) (Elurbe et al., 2017; Session et al., 2016); as expected, a greater number of peaks were identified in each of the long homeologous chromosomes than the short chromosomes (Figure 1B).

**Figure 1.**
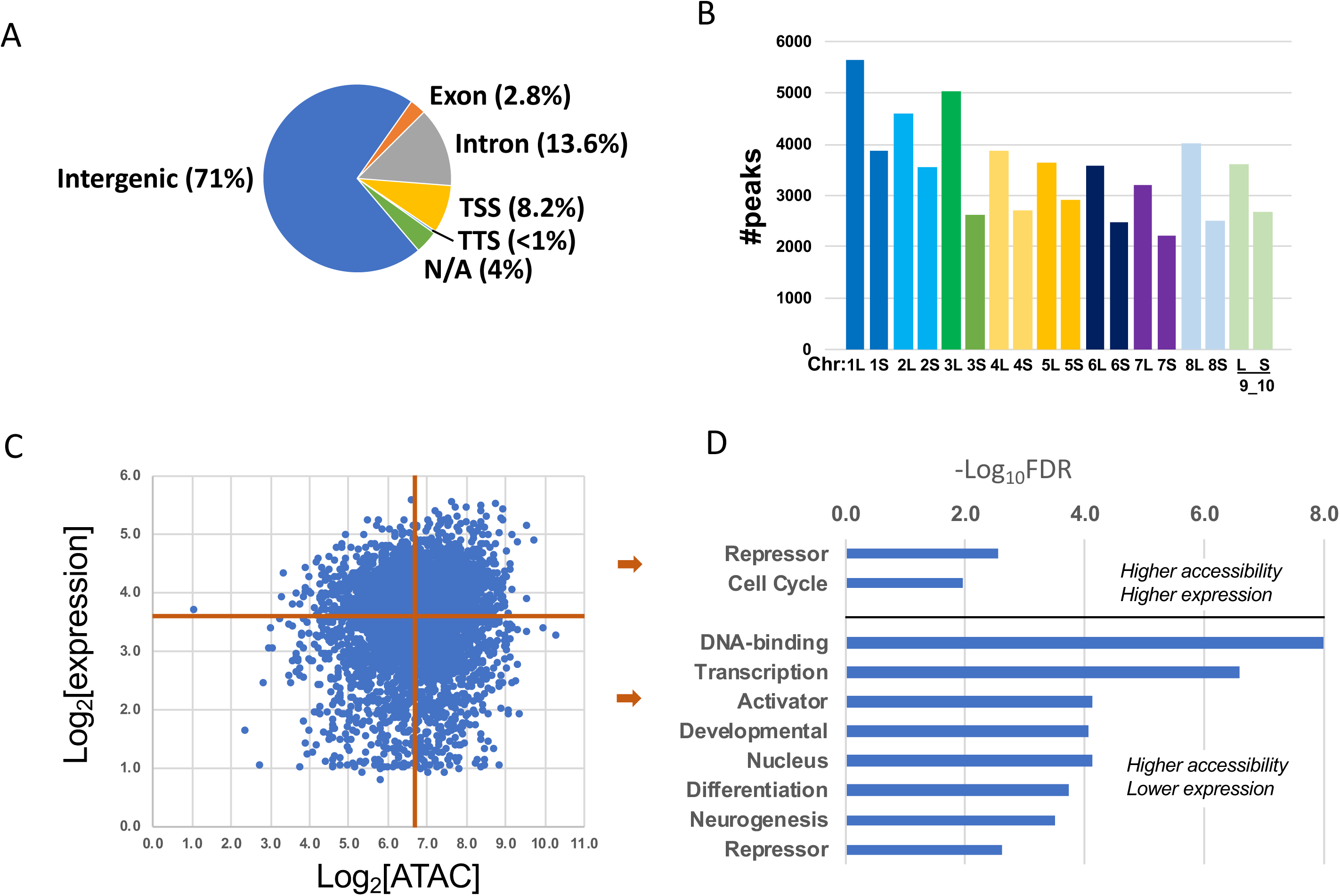
Characterization of Chromatin Accessibility at the onset of Gastrulation. **A.** Distribution of ATAC-seq peaks at the gastrula stage. ATAC-seq was performed on ectodermal explants from early gastrula (stage 10) embryos (3 biological replicates). Sequences were aligned to the xenLae2 genome and peaks were called with MACS2 and annotated with HOMER. Peaks that had 0 reads in 1 or more of the replicates (1.3% of total peaks called) were excluded from this analysis. **B.** Number of accessible regions in each chromosome. Homeologous chromosomes (“L” = long, “S”= short chromosome) are represented in the same color. An additional 6805 peaks mapped to scaffolds that have not been assigned to a chromosome. **C.** Correlation between promoter accessibility and gene expression in stage 10 ectodermal explants. Log_2_ of mRNA expression detected by microarray (Livigni et al., 2013) was plotted vs. log_2_ of normalized mean ATAC reads for peaks within 1kb of the TSS. The data were divided into 4 quadrants based on the median value for each axis so that an equal number of data points was used to analyze each quadrant. **D.** Genes from each quadrant in panel D were subjected to functional annotation using DAVID, which showed limited enrichment for terms with FDR < 0.01, except for the group with higher ATAC reads and lower expression levels (lower right quadrant in D). The −Log_10_ of the FDR is plotted on the horizontal axis.

Comparing peaks near annotated TSSs (±1kb) with published microarray data for stage 10 ectodermal explants (Livigni et al., 2013) demonstrated limited overall correlation between chromatin accessibility and gene expression (Figure 1C), as also reported previously in other cell types (Starks et al., 2019). However, segregating these genes into 4 groups based on accessibility and expression level (see (Starks et al., 2019)) revealed a striking enrichment in transcription factors in the quadrant with high accessibility and low expression (Figure 1 C,D, Supplementary Data 2), with limited enrichment in the other quadrants. These transcription factors included multiple *fox* family members (*foxa1, foxb1, foxc2, foxd1, foxd3, foxg1, foxh1, foxn4, fox01, foxo3*); *gata2,4,5,6*; *bmi1*; *irx1,2,3*; *myod; neurod; otx2; pax9, pitx2, myb, jun; sox17a, sox8*; and multiple nuclear hormone receptor family members (Supplementary Data 2).

These data present the first genome-wide characterization of chromatin accessibility using ATAC-Seq in *Xenopus* embryos and contribute to understanding the relationship between chromatin architecture and embryonic development (Akkers et al., 2009; Gupta et al., 2014; Hontelez et al., 2015; Schneider et al., 2011).

### Dorsal genes are inaccessible at the onset of gastrulation

We first examined mechanisms for loss of competence to respond to early dorsalizing signals. Induction of the dorsal organizer, one of the earliest inductive events in *Xenopus*, requires Wnt pathway activation within prospective dorsal cells during cleavage stages, well before ZGA (Blythe et al., 2010; Christian and Moon, 1993; Darken and Wilson, 2001; Heasman et al., 2000; Kao et al., 1986; Yan et al., 2018; Yang et al., 2002), and culminates with the expression of the dorsal organizer genes *sia* and *nodal3.1* after the onset of ZGA (Ding et al., 2017; Laurent et al., 1997; Lemaire et al., 1995; Smith et al., 1995). Ventral cells are also competent to respond to Wnt activation of dorsal Wnt target genes and subsequent dorsal patterning, but competence declines after the 64-cell stage and is lost by the late blastula stage (stage 9) (Figure 2A). The inability to induce dorsal Wnt target genes in the late blastula is not due to a change in overall Wnt pathway activity, however, as presumptive ventral cells are competent to respond to Wnt pathway activation by LiCl by expressing Wnt target genes involved in patterning of the anterior-posterior axis, such as *hoxa1*, *hoxd1*, and *cdx2* (Figure 2B), consistent with prior work showing that late Wnt activation causes ß-catenin stabilization, translocation to the nucleus, and binding to and activation of Wnt-target genes (Darken and Wilson, 2001; Hamilton et al., 2001; Kjolby and Harland, 2017; Nakamura et al., 2016; Schneider et al., 1996; Schohl and Fagotto, 2002).

**Figure 2.**
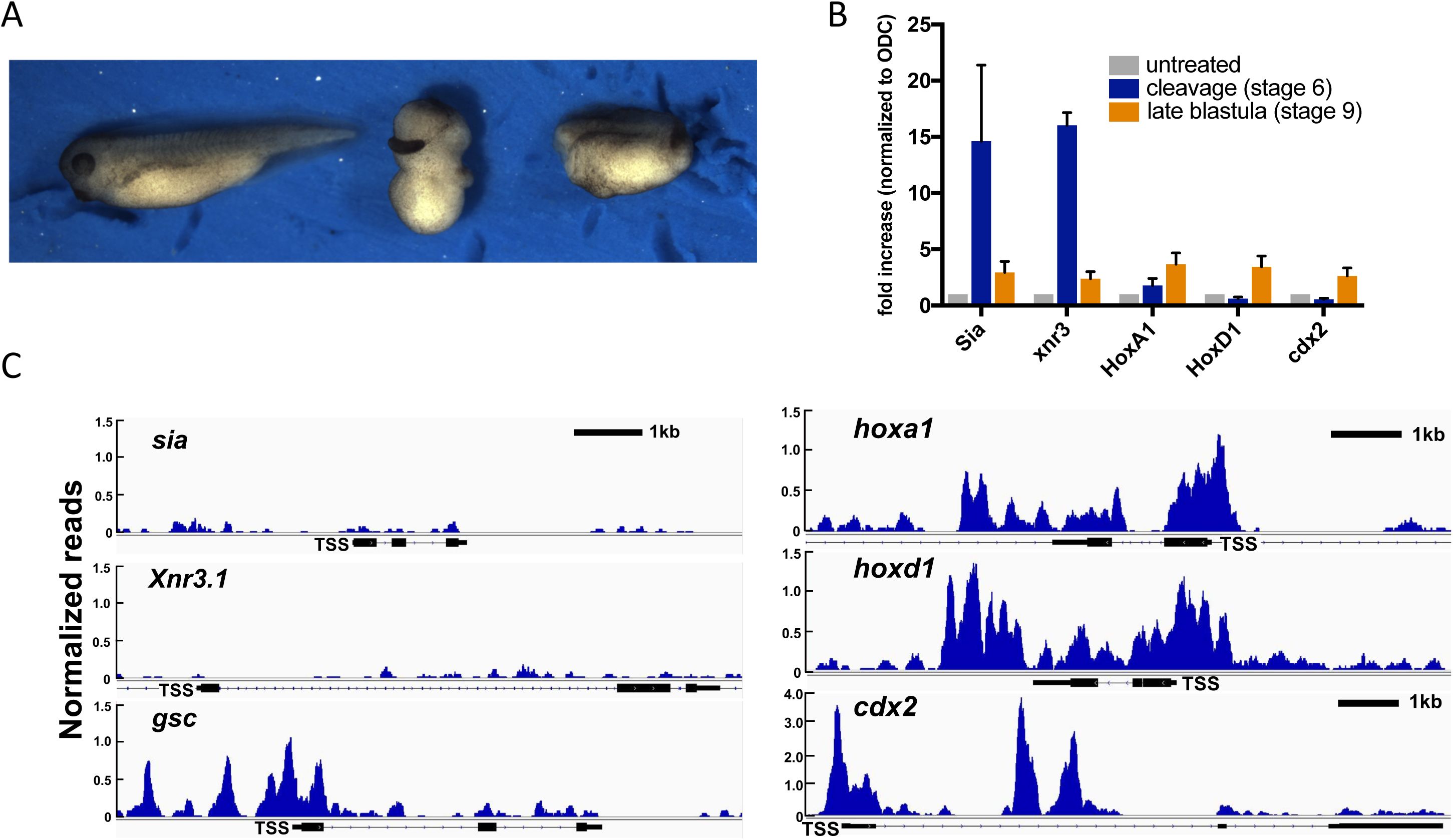
Dorsal genes are inaccessible at the onset of Gastrulation. **A.** Embryos were cultured in 0.1xMMR and Wnt signaling was activated by exposing embryos to a 10-12 minute pulse of 0.3M LiCl at the indicated stages. Activation of Wnt signaling in cleavage stage embryos leads to radial dorsalization (center). Activation of Wnt signaling at the late blastula stage leads to posteriorization (right). Controls are untreated stage 33 tadpoles. For this and following experiments, ≥ 80% of embryos treated with LiCl at the cleavage stage showed dorsalization at tadpole stages, with complete lack of trunk structures, no obvious somites, expanded and frequently circumferential eyes, and frequently expanded cement gland (DAI ≥ 8; (Kao and Elinson, 1988; Karimi et al., 2018)). Similarly, ≥80% of embryos exposed to LiCl at the late blastula stage demonstrated anterior truncation/posteriorization, with reduced or absent forebrain and small or absent cement gland, frequently with small or absent eyes and microcephaly. **B.** Wnt signaling was activated by exposure to LiCl at cleavage stage (stage 6) or the blastula stage (stage 9) and expression of dorsal Wnt target genes *sia1* and *nodal3.1* and non-dorsal Wnt target genes (*hoxa1, hoxd1,* and *cdx2*) was measured by RT-qPCR in whole embryos at the midgastrula stage (stage 10.5). Gene expression in all samples was normalized to *ODC* and then expression in embryos exposed to LiCl at stage 6 (blue bars) and stage 9 (orange bars) was normalized to untreated (grey) and are presented as mean values for fold change in expression for 5 biological replicates except for *nodal3.1* which had 3 replicates. Error bars represent standard error of the mean (SEM). **C.** Representative sequencing tracks for the dorsal Wnt target genes *sia1* and *nodal3.1*, the organizer marker *Gsc,* and non-dorsal Wnt target genes *hoxa1*, *hoxd1*, and *cdx2*. Y-axis represents normalized reads scaled by 1,000,000/total alignments. Scale bar represents 1kb.

As the Wnt signaling pathway is active in presumptive ventral cells of the late blastula (when competence to induce dorsal genes is lost) we hypothesize that loss of competence is associated with inaccessibility of regulatory regions at dorsal Wnt target gene promoters.

Consistent with this hypothesis, the ATAC-seq data show that promoters of the dorsal Wnt target genes *sia1, sia2,* and *nodal3.1* are not accessible at the early gastrula stage whereas the Wnt-inducible genes *hoxa1*, *hoxd1*, as well as the canonical Wnt target gene *axin2* are accessible (Figure 2E, Supplementary Data 1). Thus, loss of competence to Wnt signaling to induce dorsal genes correlates with inaccessibility of dorsal Wnt target gene promoters.

### Repressive histone modifications are not associated with loss of competence to respond to early dorsalizing signals

The lack of accessibility at the *sia1* and *nodal3.1* promoters suggests that the chromatin at these sites is in a repressed state. As chromatin inaccessibility and transcriptional repression are associated with specific histone modifications, such as trimethylation of histone H3 at lysine-9 (H3K9me3) and lysine-27 (H3K27me3) (Perino and Veenstra, 2016), we hypothesized that deposition of these repressive marks would be associated with loss of competence. Specifically, we predicted that repressive histone modifications would preferentially accumulate in the ventral cells at the *sia1* and *nodal3.1* promoters (Figure 3A). To test this, we dissected early gastrula (stage 10) embryos into dorsal and ventral halves and then performed chromatin immunoprecipitation (ChIP) followed by qPCR for the *sia1* and *nodal3.1* promoters (Blythe et al., 2010) to compare the abundance of H3K27me3 and H3K9me3 in ventral versus dorsal cells. H3K27me3 was undetectable at the *sia1* and *nodal3.1* promoters in ventral and dorsal halves of early gastrula embryos (Figure 3B). We were, however, able to detect H3K27me3 at the *gsc* promoter in gastrula stage embryos (Figure 3C). The low abundance of repressive marks at these promoters is consistent with ChIP-Seq data from the closely related species *X. tropicalis* that show H3K27me3 begins to accumulate at spatially restricted, developmentally regulated genes during the mid to late gastrula stage (stage 11 and later) (Akkers et al., 2009). The appearance of H3K27me3 during the late gastrula, long after loss of competence, suggests H3K27me3 is not a feature of loss of competence to Wnt during dorsal induction. Furthermore, H3K9me3 at *sia1* and *nodal3.1* was barely detectable above background in dorsal and ventral cells compared to control genes *xretpos(L)* and *1a11*, retrotransposons that have acquired H3K9me3 by the gastrula stage (Herberg et al., 2015), and there was no detectable enrichment in ventral halves (Figure 3D). These findings do not support a role for H3K9me3 or H3K27me3 in the loss of competence to respond to Wnt signaling at *sia1* or *nodal3.1*.

**Figure 3.**
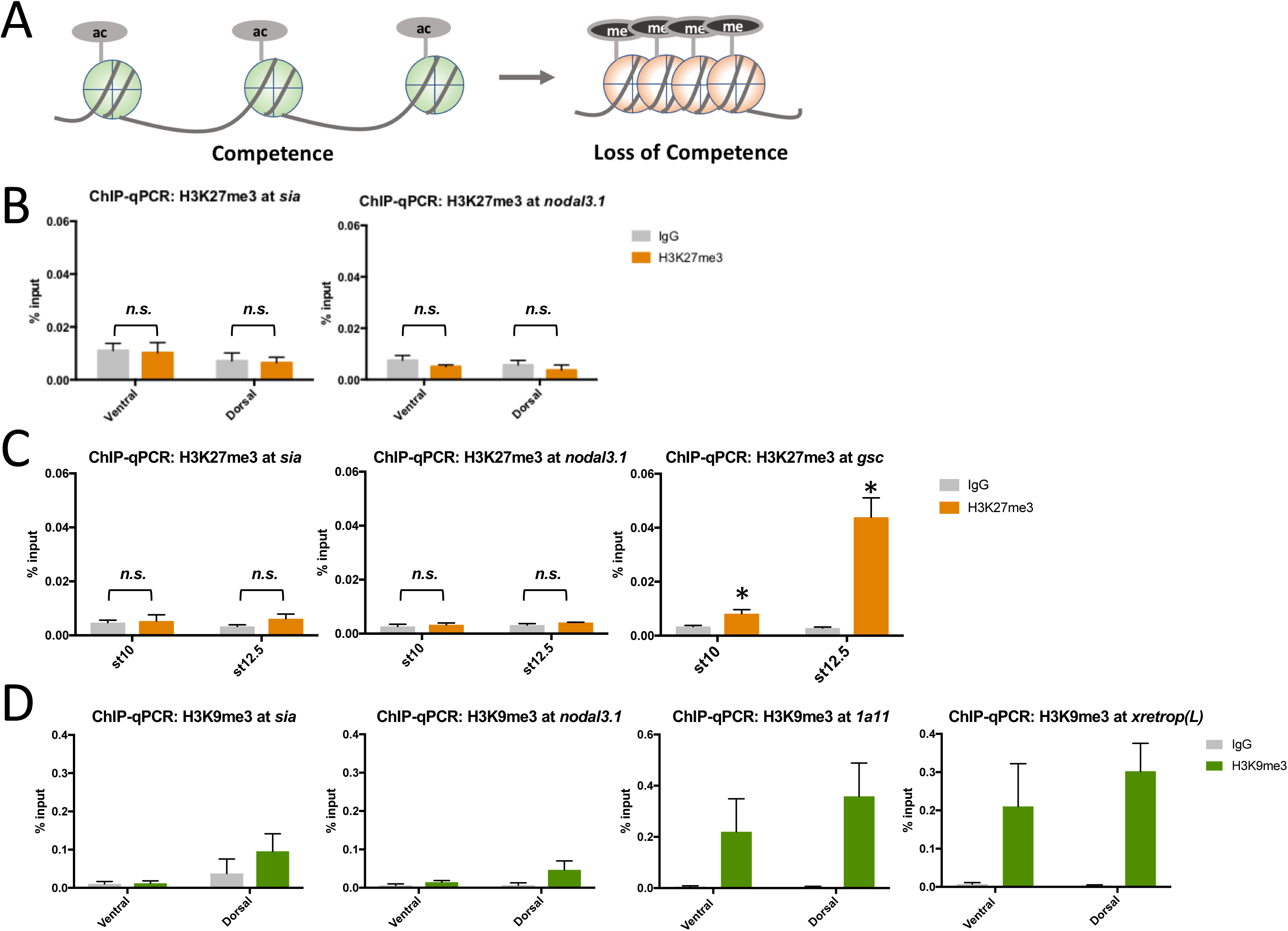
Repressive histone modifications H3K9me3 and H3K27me3 are not associated with loss of competence. **A.** Model for chromatin accessibility mediating loss of competence. Left: Open chromatin state predicted to be associated with competence. “Ac” represents histone acetylation, typically associated with active or accessible chromatin. Right: Chromatin inaccessibility predicted to be associated with loss of competence. “Me” represents histone methylation at sites that are typically associated with repressive states or closed chromatin, such as H3K9me3 or H3K27me. **B**. ChIP-qPCR for H3K27me3 (orange) at Wnt target genes *sia1* and *nodal3.1* in dorsal versus ventral halves of gastrula stage (stage 10) embryos. IgG controls in grey. Embryos were cultured in 0.1xMMR until gastrula stage, fixed, dissected into dorsal and ventral halves using the dorsal blastopore lip as a marker, and subjected to ChIP. **C.** ChIP-qPCR for H3K27me3 in whole embryos at the *sia1, nodal3.1,* and *gsc* promoters was also performed at early (st 10) and late (st 12.5) gastrula stages. \**p* < 0.05 for comparison of H3K27me3 to IgG background control (one-tailed t-test); “*n.s.*” indicates not significant. **D.** ChIP-qPCR for H3K9me3 (green) at promoters for *sia1* and *nodal3.1,* as well the control genes *1a11* and *xretrop(L),* in dorsal and ventral halves of gastrula stage embryos. IgG controls in grey. Embryos cultured, dissected, and collected as in B. Data in B - D represent means of 3 biological replicates and error bars show SEM.

### Inhibition of histone deacetylases extends competence to early dorsalizing signals

Because of technical limitiations of ATAC-Seq in *Xenopus laevis* embryos, we were unable to assess chromatin accessibility at the time of maximal competence to respond to dorsal Wnt signals (32-64 cell stage) or at any time prior to the late blastula stage. However, open chromatin and active transcription tend to be associated with histone acetylation, including lysine 9 and 14 of histone H3 (H3K9 and H3K14) near promoters and lysine 27 of histone H3 (H3K27) at active enhancers (Shahbazian and Grunstein, 2007). Wnt signaling promotes acetylation of target gene promoters by stabilizing ß-catenin, which binds to the transcription factor TCF and recruits the histone acetyltransferase p300 (Hecht et al., 2000). In the absence of Wnt signaling, TCF proteins repress target gene expression by recruiting histone deacetylases (HDACs) through diverse corepressors (Cadigan, 2012; Ramakrishnan et al., 2018). Therefore, in regions of the embryo that are competent to respond but do not receive a Wnt signal, such as ventral blastomeres in the cleavage stage embryo, dorsal Wnt target gene promoters may become deacetylated and inaccessible, as we observed for *sia1* and *nodal3.1* at the early gastrula stage. Prior work had shown that the *sia1* and *nodal3.1* promoters are robustly acetylated at H3K9/K14 prior to the midblastula transition (MBT) and are associated with the activating histone mark H3K4me3 and the initiating form of RNA polymerase II (Blythe et al., 2010). However, at the late blastula stage, H3K9 acetylation is barely detectable at the *sia1* and *nodal3.1* promoters (Figure 4A). To test whether histone deacetylation contributes to loss of competence, we inhibited HDACs with Trichostatin-A (TSA) beginning at the 64-cell stage and then assessed histone acetylation and Wnt responsiveness at the late blastula stage. TSA caused a marked increase in H3K9 acetylation at the *sia1* and *nodal3.1* promoters compared to untreated embryos as measured by CHIP-qPCR. (Figure 4A). This response to HDAC inhibition indicates that acetylation at dorsal Wnt target gene promoters is dynamic, with acetylation continuing when HDACs are inhibited, allowing us to test whether acetylation at dorsal Wnt target genes would maintain competence at the late blastula stage.

**Figure 4.**
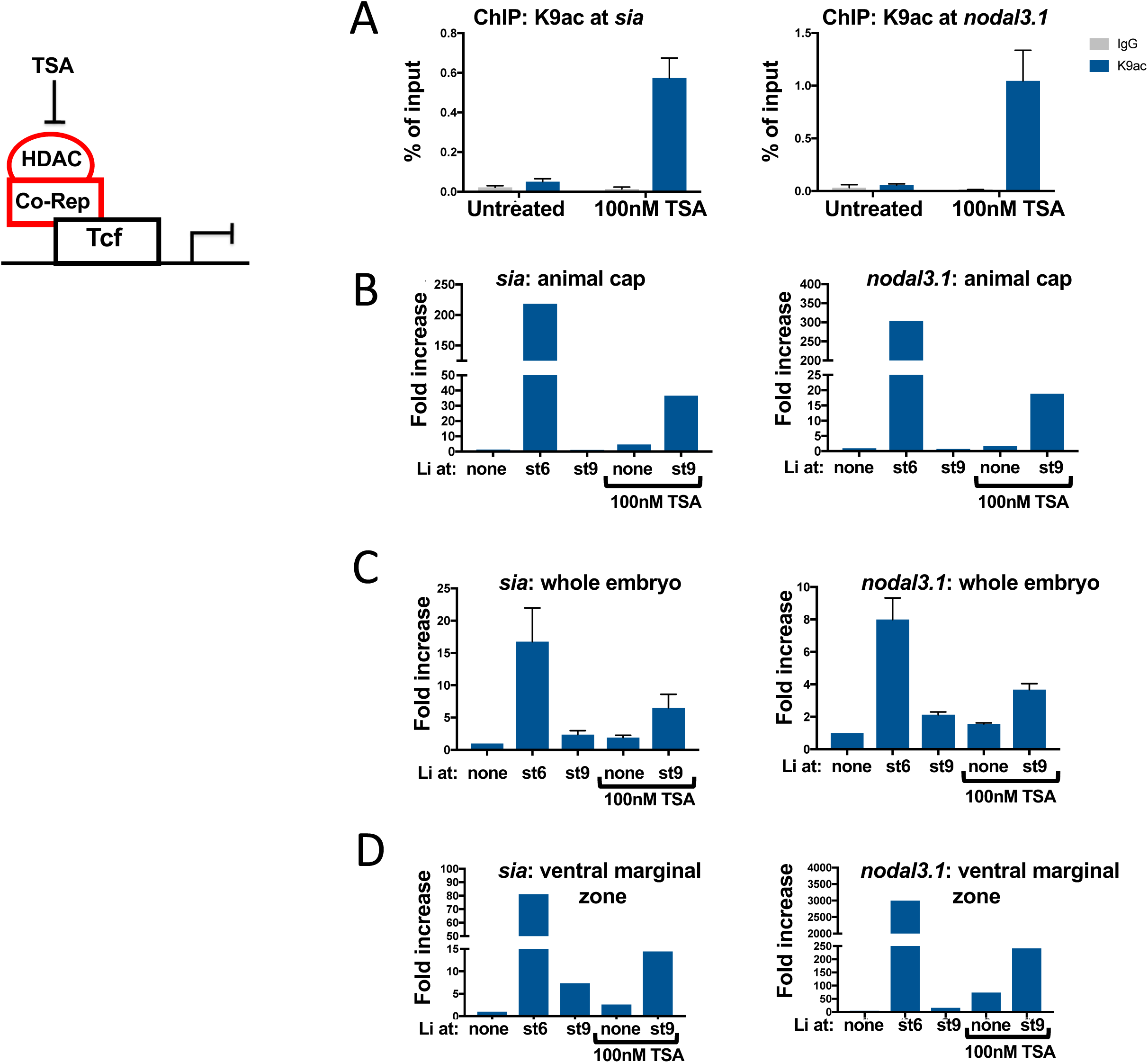
Histone deacetylase activity suppresses competence for dorsal induction. **A.** Embryos were cultured with and without 100 nM Trichostatin A (TSA) beginning at the 64-cell stage and collected at the onset of gastrulation. ChIP-qPCR was performed for H3K9ac (blue) at the *<u>sia1</u>* and *nodal3.1* promoters. IgG controls are in grey. **B.** Embryos were treated with or without 100 nM TSA beginning at the 32-64 cell stage and exposed to LiCl in the late blastula stage as in Figure 2. Ectodermal explants were dissected at the onset of gastrulation and harvested immediately to measure expression of *sia1* and *nodal3.1* as in Figure 2. As *sia1* and *nodal3.1* were frequently undetectable in explants from control embryos, it was not always possible to calculate a fold-increase relative to control. The histogram shows the mean of two replicates for which *sia1* was detectable in controls. Regression analysis found a significant increase in both *sia1* and *nodal3.1* expression in explants treated at stage 9 with TSA and lithium compared to lithium alone (for *sia1*, mean ΔΔCt = 2.09 with p < 0.0001; for *nodal3.1*, mean ΔΔCt = 2.65, p < 0.0001) or TSA alone (for *sia1*, mean ΔΔCt = 1.67 with p = 0.0004; for *nodal3.1*, mean ΔΔCt = 1.65 with p = 0.008); there was no significant interaction between lithium and TSA. **C.** Embryos were exposed to LiCl and TSA as in panel B and harvested at the onset of gastrulation to measure whole embryo expression of *sia1* and *nodal3.1*. **D.** Embryos were exposed to LiCl and TSA as above and ventral marginal zones were dissected at the onset of gastrulation and harvested immediately to measure expression of *sia1* and *nodal3.1*.

To test whether histone deacetylation contributes to loss of competence, we first used ectodermal explants (animal caps), which do not normally express *sia1* or *nodal3.1* but are competent to express these dorsal genes in response to Wnt pathway activation during early cleavage stages and lose competence for dorsal gene induction after the MBT, similar to whole embryos (Carnac et al., 1996; Christian and Moon, 1993; Fagotto et al., 1997; Yang-Snyder et al., 1996). Embryos were cultured in control medium or medium containing TSA (beginning at the 64-cell stage (Stage 6)) and then exposed to control buffer or a pulse of lithium chloride (LiCl) at the late blastula stage (Stage 9) to activate Wnt signaling. As a positive control, a separate group of embryos was treated with a pulse of LiCl at stage 6. At the onset of gastrulation, explants were dissected and harvested. Expression of *sia1* and *nodal3.1* was then measured by reverse-transcription-quantitative PCR (RT-qPCR). Expression of *sia1* and *nodal3.1* in untreated control explants was frequently undetectable, making it difficult to report fold-change in expression relative to control. We addressed this in two ways: we performed the experiment with 10 independent clutches of embryos and Figure 4B shows the mean of experiments in which we could report fold change in dorsal gene expression relative to control explants. In addition, to compare the effect of lithium and TSA treatment at stage 9 in all 10 replicates, we used generalized linear models fitted with generalized estimating equations (GEE) to compare ΔCt values, as described in detail in methods. Regression results revealed a significant induction of both *sia1* and *nodal3.1* by lithium at stage 9 in TSA-treated embryos compared to lithium or TSA alone (see legend to Figure 4B). Interaction terms were not significant, and were omitted from the final models. These observations support the hypothesis that histone deacetylation contributes to the loss of competence in presumptive ectoderm.

To confirm HDAC regulation of Wnt-target gene competence in whole embryos, embryos were treated with TSA beginning at the 64-cell stage, exposed to LiCl in the late blastula stage (stage 9), and *sia1* and *nodal3.1* expression was measured at the onset of gastrulation. As in the ectodermal explants, HDAC inhibition maintained competence in intact embryos such that *sia1* and *nodal3.1* were induced 10-fold and 4-fold, respectively, in response to Wnt signaling at the late blastula stage (Figure 4C). To ensure that HDAC inhibition maintains competence specifically in ventral cells, we treated whole embryos with TSA and dissected ventral marginal zones (VMZs) at the onset of gastrulation. Expression of *sia1* and *nodal3.1* was low in control VMZs and VMZs from embryos exposed to LiCl at the blastula stage. In contrast, LiCl treatment at the late blastula stage in embryos treated with TSA induced dorsal Wnt target genes in VMZs to a greater extent than LiCl or TSA alone. Thus, similar to ectodermal explants and whole embryos, inhibition of HDAC activity maintains competence of ventral cells to express *sia1* and *nodal3.1* in response to Wnt signaling at the late blastula stage (Figure 4D). That chromatin at *sia1* and *nodal3.1* is inaccessible at gastrulation yet HDAC inhibition increases acetylation at these promoters and extends competence to Wnt activation in late bastula stage embryos suggests that deacetylation of histones at Wnt target gene promoters contributes to the loss of competence to respond to Wnt signaling.

### Chromatin accessibility and histone deacetylation do not correlate with loss of competence for mesoderm induction by FGF

We also asked whether closed chromatin and histone deacetylation are common features of loss of competence during later developmental stages. Classic recombinant experiments in amphibian embryos demonstrated that a signal from the vegetal hemisphere of blastula stage embryos induces mesoderm in animal hemisphere cells fated to become ectoderm (Nakamura et al., 1970; Sudarwati and Nieuwkoop, 1971) and in *Xenopus* the competence of ectoderm to respond to this inducing signal is lost at the gastrula stage (Dale et al., 1985; Gurdon et al., 1985; Jones and Woodland, 1987). Similarly, fibroblast growth factor (FGF) and TGF-ß family ligands mimic the vegetal mesoderm inducing signal and competence of ectoderm to respond to FGF is lost at the gastrula stage (Asashima et al., 1990; Green et al., 1990; Rosa et al., 1988; Slack et al., 1987; Sokol et al., 1990). To ask whether the loss of accessibility at the promoters for mesodermal genes correlates with loss of competence to respond to FGF, we performed ATAC-seq in late gastrula (stage 12) ectodermal explants and compared accessibility in early gastrulae (stage 10), when explants are competent to respond to FGF. ATAC-seq data from early gastrula stage ectoderm demonstrated that chromatin remains open at the promoters for mesodermal genes (Figure 5A, Supplementary Data 1), including *myod1 (myoD)*, *wnt8a,* and *tbxt* (*brachyury/Xbra*), a master regulator of mesoderm induction that is strongly induced by FGF during the blastula stage but no longer competent to respond to FGF at the gastrula stage (stage 10) (Slack et al., 1987). Furthermore, the promoters for *tbxt, myod1*, and *wnt8a* remain open through stage 12 (Supplementary Data 1, 3). Based on these observations, we predicted that mesoderm induction, and specifically induction of *tbxt*, may not be sensitive to HDAC inhibition. However, it remained possible that deacetylation at regulatory sites apart from the promoter may be responsible for the loss of competence to mesoderm inducing signals. We therefore inhibited HDACs with TSA beginning at late blastula (stage 9) and then dissected and cultured ectodermal explants with FGF at the onset of gastrulation, when control explants lose competence (Figure 5B). Explants were then collected at stage 13 to assess *tbxt* expression. FGF robustly induced *tbxt* expression in explants treated at the blastula stage but the response at the gastrula stage was markedly reduced (as expected) and was not enhanced or maintained by HDAC inhibition (Figure 5C).

**Figure 5.**
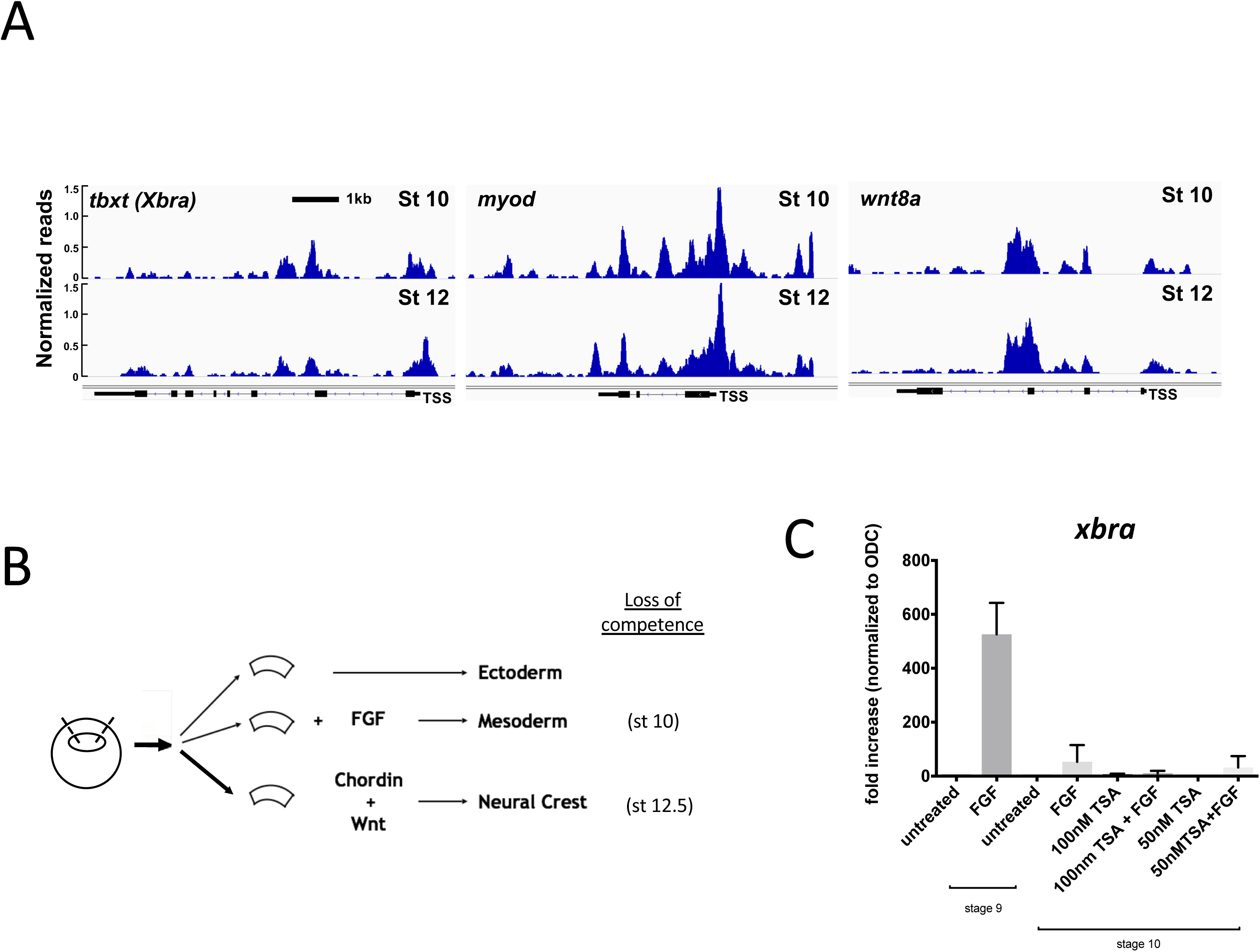
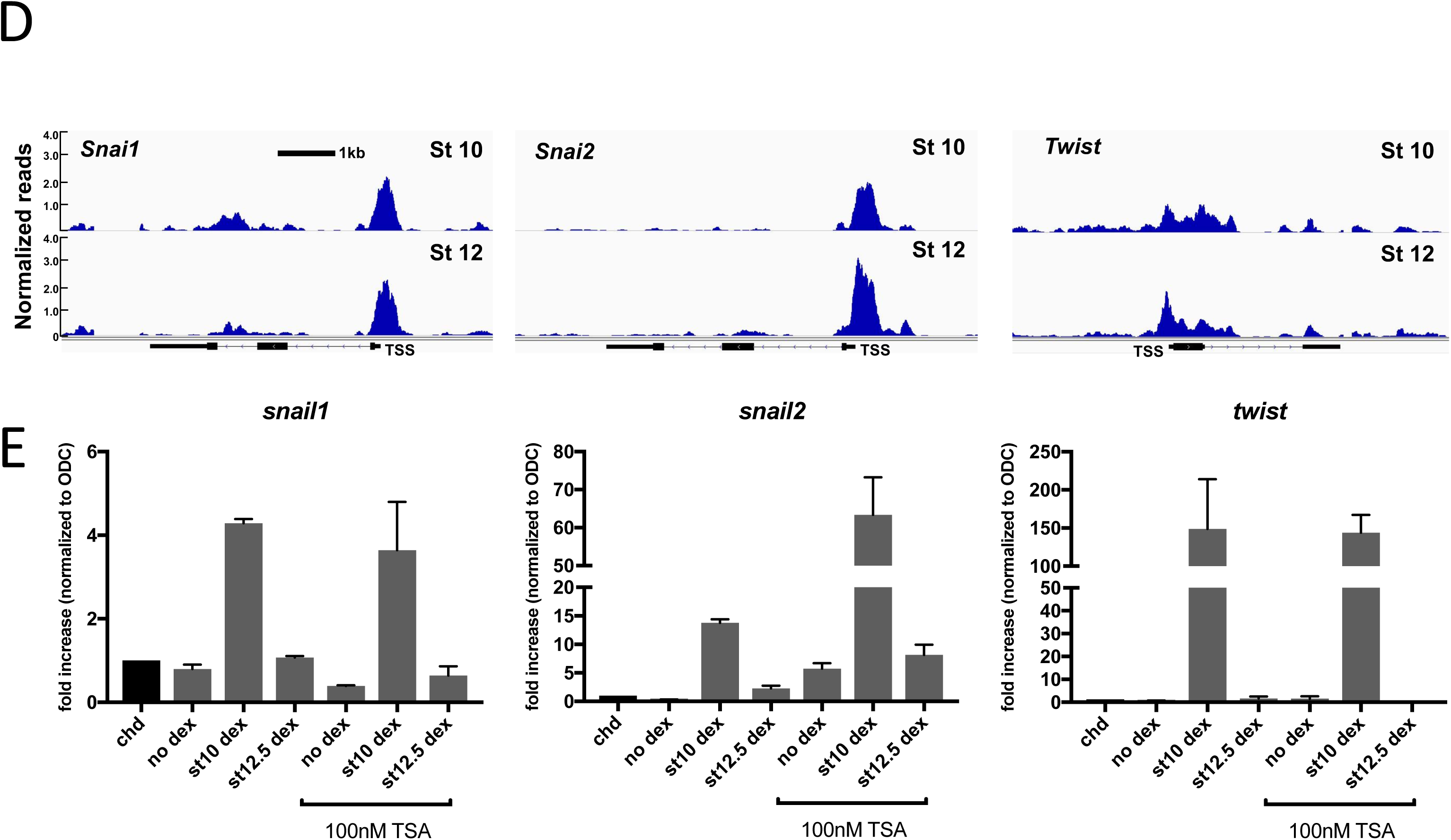
Chromatin Accessibility and histone deacetylation do not correlate with loss of competence at later stages of development. **A.** Representative sequencing tracks for the mesoderm genes *tbxt*, *wnt8a*, and *myod1* at stage 10. The ATAC-seq data are normalized to total reads. Y-axis represents the number of normalized reads with the scale chosen for optimal visualization of peaks for each gene. Scale bar represents 1kb in all panels. **B.** Diagram of explant assays for mesoderm and neural crest induction. On the left is a blastula embryo showing where ectodermal explants are dissected (dashed lines). In the center are explants that can be subjected to different treatments (FGF or Wnts) to induce mesoderm or neural crest, respectively. On the right are the stages at which the ectodermal explant loses competence for a given inductive signal. **C.** For mesoderm induction, explants were dissected at the late blastula stage and cultured in FGF or control buffer and harvested at stage 14 for analysis of *tbxt* expression by RT-qPCR. A separate group of embryos was treated with or without 100nM TSA beginning at the late blastula stage (stage 9) until ectodermal explants were dissected at the gastrula stage. Explants were similarly cultured with or without FGF at early gastrula (stage 10.5) and harvested at stage 14 to measure induction of *tbxt*. All groups are normalized to untreated controls. **D.** Representative sequencing tracks for neural crest genes (*snai1*, *snai2*, *twist*) during competence (stage 10) and at loss of competence (stage 12). The ATAC-seq data are normalized to total reads. Y-axis represents the number of normalized reads with the scale chosen for optimal visualization of peaks for each gene. Scale bar represents 1kb. **E.** RT-qPCR measuring gene expression of neural crest genes (*snai1*, *snai2*, *twist*) normalized to housekeeping gene *ODC*. Each blastomere of 2-cell embryos was injected in the animal pole with mRNAs encoding Chordin (0.5ng/blastomere) and THVGR (10pg/blastomere). Ectodermal explants were dissected at the late blastula stage (stage 9) and cultured until early gastrula (stage 10). A subset of explants were cultured in control buffer and then treated with dexamethasone at early gastrula stage (stage 10) to induce THVGR and activate Wnt signaling. The rest of the ectodermal explants were cultured in 100nm TSA. Of the TSA treated explants, a subset was treated with dexamethasone at early gastrula (stage 10) or at late gastrula (stage 12.5). Explants were cultured until stage 22 when they were collected and expression for *snai1*, *snai2*, *twist* was measured by RT-qPCR. Data in panels C and E represent 3 biological replicates and error bars are SEM.

### Chromatin accessibility and histone deacetylation do not correlate with loss of competence to Wnt in neural crest induction

While loss of competence for mesoderm induction is not associated with loss of chromatin accessibility at multiple mesodermal gene promoters, histone deacetylation could still be a general mechanism for loss of competence to respond to Wnts, which act iteratively throughout early development. For example, Wnt signaling is required for neural crest induction at the border of the neural plate and non-neural ectoderm (Aybar and Mayor, 2002; Saint-Jeannet et al., 1997; Wu et al., 2003). Neural ectoderm is competent to form neural crest in response to Wnts during early gastrulation but competence declines during gastrulation (stage 12) and is lost by the beginning of the neurula stage (Mancilla and Mayor, 1996). Early markers and essential regulators of neural crest induction include the direct Wnt target genes *snai1* (*snail/snail1)*, *snai2* (formerly *slug/snail2)*, and *twist* (Aybar and Mayor, 2002; Wu et al., 2003). Similar to mesodermal gene promoters, and in contrast to dorsal Wnt target genes, the chromatin at promoters of *snai1*, *snai2*, and *twist* remained open at both early and late gastrula stages (Figure 5D, Supplementary Data 1), as well as at the neurula stage (data not shown), arguing against a loss of chromatin accessibility at these promoters as an explanation for the loss of response. Similarly, inhibiting HDACs did not maintain competence to induce neural crest (Figure 5B). Thus, using a well-established neural crest induction assay (Saint-Jeannet et al., 1997), we dissected ectodermal explants from embryos expressing *chrd*, activated Wnt signaling at either stage 10 or stage 12.5 using a glucocorticoid-inducible activated version of TCF (THVGR (Darken and Wilson, 2001; Wu et al., 2005; Yang et al., 2002)), and harvested explants at the tailbud stage (stage 22). Activation of THVGR at stage 10 strongly induced *snai1*, *snai2*, and *twist* whereas there was minimal induction when THVGR was activated at stage 12.5, consistent with previously established competence windows (Mancilla and Mayor, 1996). Unlike dorsal competence, HDAC inhibition (by addition of TSA at the time of explant dissections) did not extend competence to induce neural crest genes by Wnt signaling (Figure 5E). Higher concentrations (e.g. 200nM) or longer exposure to TSA blocked the induction of neural crest markers by THVGR (consistent with a report that HDAC activity is required for neural crest induction (Rao and LaBonne, 2018)) and precluded us from further investigating the role of histone deacetylation in competence during neural crest induction (data not shown). These results suggest that the mechanisms for loss of competence to Wnt signaling are context dependent.

### Differential Chromatin Accessibility at early versus late gastrulation

Although mesodermal genes, such as *tbxt*, *myod1*, and *wnt8a,* and Wnt-inducible neural crest genes including *snai1*, *snai2*, and *twist* retain open chromatin at their promoters after loss of competence, changes in chromatin accessibility at other sites involved in these inductive processes could instead be associated with loss of competence. To understand more about the potential role of loss of chromatin accessibility in modulating the response to inductive signals, we took three approaches.

First, we compared genome-wide chromatin accessibility in stage 10 versus stage 12 ectodermal explants. There were ∼70,000 called peaks at stage 12, similar to stage 10. Of these, 9070 peaks were located within 1kb of annotated TSSs at stage 10 and 9108 at stage 12 and overall there was a strong correlation in peak intensity between stages 10 and 12 (Figure 6A, Supplementary Data 1). We also compared ATAC-Seq tracks for multiple *X. laevis* genes at the gastrula stage to recently reported Dnase-Seq data from stage 8.5 *X. tropicalis* embryos (Gentsch et al., 2019) and observed remarkably similar patterns of accessibility at the promoters and within the gene bodies for *tbxt, chrd, vegt, hoxa1, hoxd1, cdx2, snai1, snai2, hesx1, foxh1, wnt8a*, and other genes (Supplementary Figure 1 and data not shown), raising the possibility that patterns of accessibility established by the MBT are maintained through the gastrula stage and may be conserved between *X. laevis* and *X. tropicalis*. However, accessibility declined ≥ 2-fold at stage 12 at promoters for 279 genes and increased at stage 12 for 264 genes (Figure 6B, Supplementary Data 1). Although there was no enrichment of GO terms for genes with increased promoter accessibility at stage 12, the set of promoters with reduced accessibility at stage 12 was highly enriched for genes that regulate development, primarily transcription factors and especially homeodomain-containing factors (Figure 6C, Supplementary Data 1). Multiple genes that regulate mesodermal and neural development and patterning were enriched in this group, including *chordin*, *vegt*, *pygo2, pax3, neurogenin3*, and the pluripotency factors *pou3f1 (oct6), pou3f2,* and *pou5f3 (oct91)* (Figure 6D, 7C, Supplementary Data 1). Thus, loss of chromatin accessibility at these promoters could play a role in the loss of competence during the gastrula stage.

**Figure 6.**
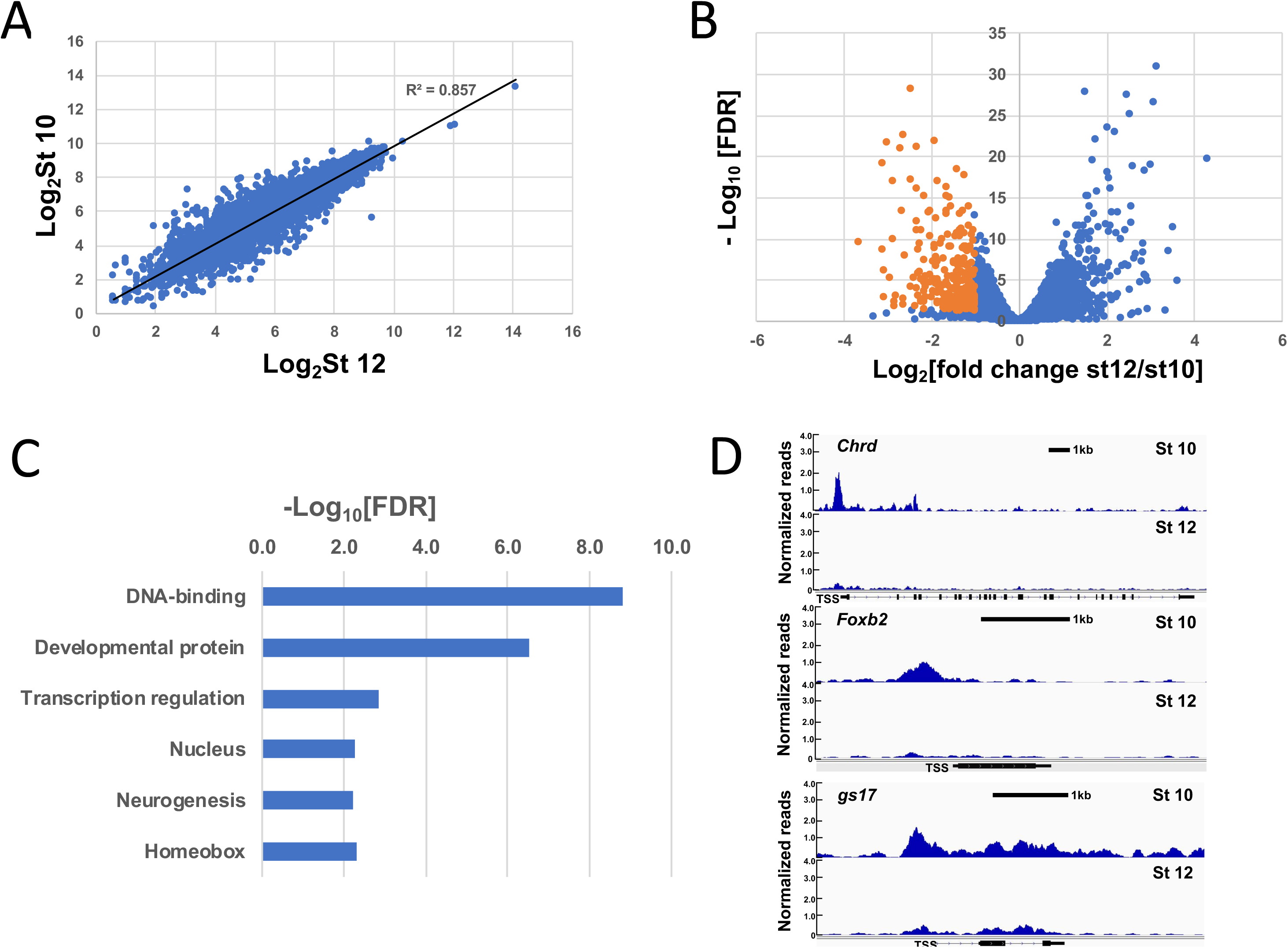
Differential Chromatin Accessibility at early versus late gastrulation. **A.** Correlation of ATAC peaks within 1kb of TSS at stage 10 vs stage 12 (FDR < 0.05). **B.** Volcano plot of Log_2_ fold change in ATAC promoter peaks at stage 12/stage 10 versus −Log_10_ of false discovery rate (FDR). Dotted lines indicate the threshold for genes selected for functional annotation. **C.** Genes associated with ATAC peaks within 1kb of the TSS that decrease by ≥ 2 fold at stage 12 were submitted for functional annotation through DAVID, which revealed marked enrichment of DNA-templated transcription factors. Uniprot keywords are ranked according to false discovery rate with a cutoff FDR < 0.01. The horizontal axis shows −Log_10_ of the FDR. A similar analysis of promoter peaks that increased ≥ 2 fold did not identify any enriched categories. **D.** Representative tracks for *chordin (chrd), foxb2*, and the temporal marker *gs17* showing dynamic chromatin accessibility at the respective promoters.

Second we examined the correlation between accessibility and p300 binding, which is associated with both poised and active cis-regulatory elements. ChIP-Seq for p300 at the gastrula stage was reported previously but was mapped to an earlier genome build (Session et al., 2016). We re-mapped the raw sequences (GSE76059) to the xenlae2 genome and identified 33,289 p300 peaks, of which 22,440 (67%) overlapped with ATAC-Seq peaks at stage 10 (Figure 7A, Supplementary Data 1, Supplementary Data 3). Of the 9252 intergenic and intronic regions that lose accessibility between stage 10 and stage 12, 2743 were associated with p300 binding, suggesting that these ATAC-accessible, p300-bound peaks represent putative cis-regulatory modules (pCRMs) that are inactivated during gastrulation. We then selected those genes with TSSs within 100kb of a pCRM and subjected them to functional gene annotation using DAVID, which revealed enrichment of terms including developmental proteins, neurogenesis, transcription regulation, homeobox, and gastrulation (Figure 7B, Supplementary Data 3). Furthermore, nearly half of the genes that lose accessibility at their promoters (Figure 6B, C) also lose accessibility at promoter-distal pCRMs, including *foxh1*, *vegt, lefty, sox17b, pax3, pou3f2,* and *wnt5b* (Figure 7C, Supplementary Data 3, and data not shown). Furthermore, the loss of accessibility at both promoters and promoter-distal pCRMs for a set of genes enriched in developmental regulators is consistent with the hypothesis that loss of chromatin accessibility may be an important mechanism for restricting developmental competence.

**Figure 7.**
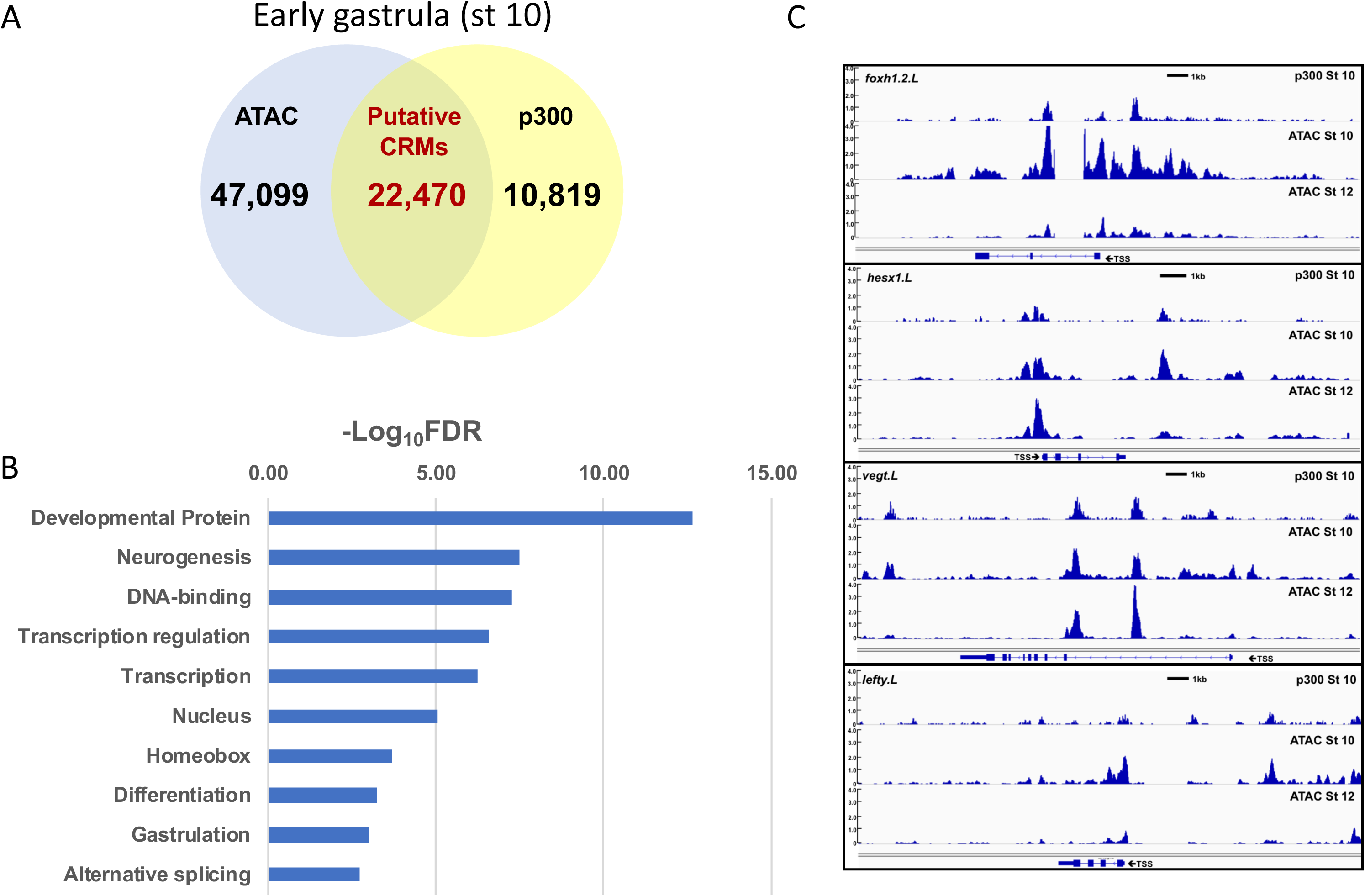
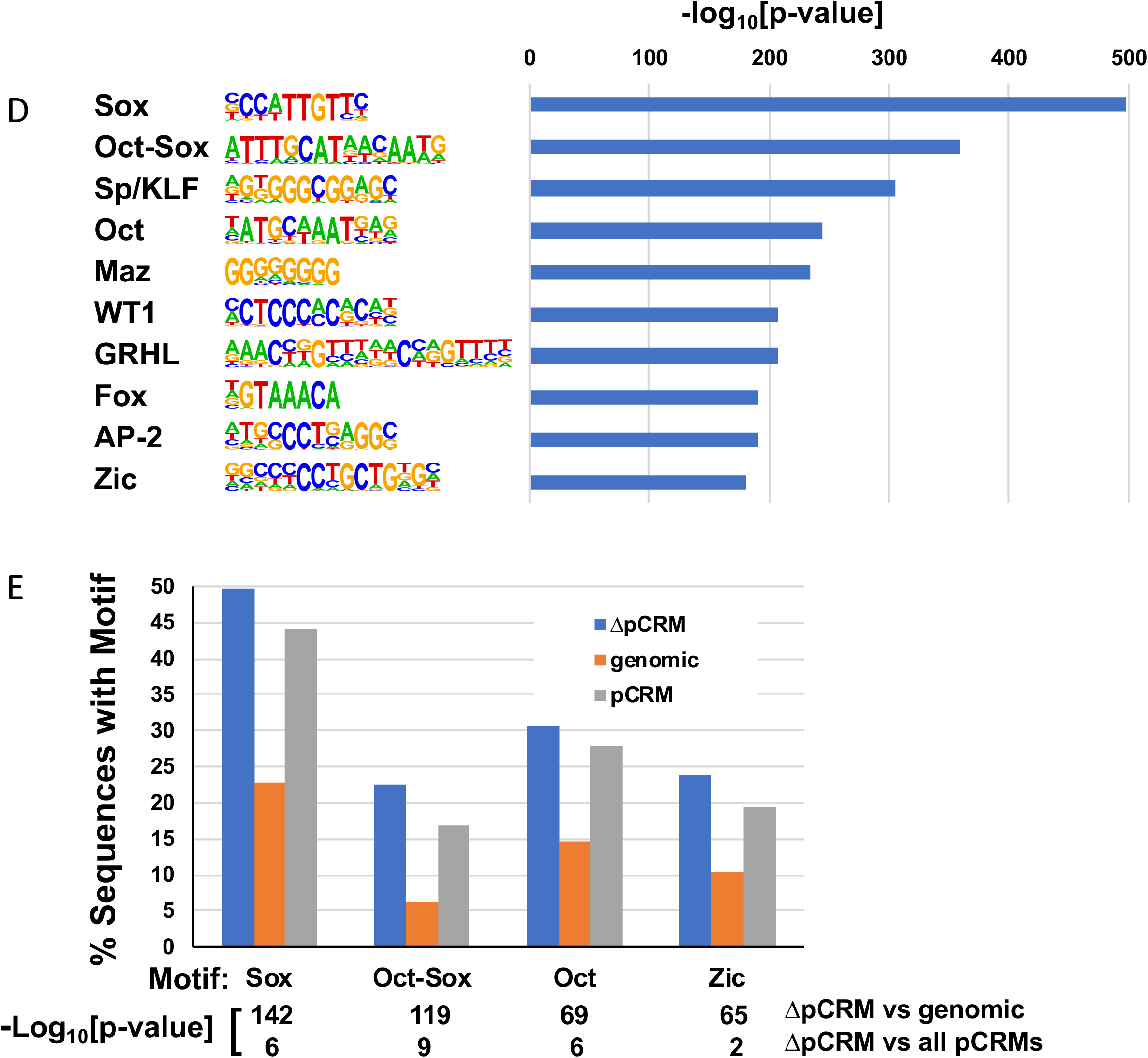
Putative Cis-regulatory Modules in gastrula stage embryos. **A.** Venn diagram of accessible regions at stage 10 (blue circle) versus p300 ChIP-seq peaks (yellow circle); overlap represents putative cis-regulatory elements. p300 peaks were determined by re-mapping ChIP-Seq data from (Session et al., 2016) to the xenlae2 genome. **B.** Accessible peaks within intergenic and intronic regions that were also bound by p300 (pCRMs) at stage 10 and which showed reduced accessibility at stage 12 were identified; genes with TSSs within 100kb of these peaks (2074) were subjected to functional gene annotation using DAVID. Uniprot keywords are ranked according to false discovery rate with a cutoff FDR < 0.01. Representative tracks for *foxh1.2, hesx1, vegt*, and *lefty* comparing p300-bound sites (top) to ATAC-Seq peaks at stage 10 (middle) and stage 12 (lower tracks). Scale bar = 1kb. **D.** Transcription factor motifs associated with stage 10 pCRMs identified by HOMER (Heinz et al., 2010). Sites are ranked by −log10[p-value] (compared to genomic background); those with p ≤ 10^-170^ are shown. **E.** Frequency of transcription factor motifs in pCRMs (blue) that showed reduced accessibility at stage 12 (ΔpCRMs) compared to genomic background (orange) and to all pCRMs (gray). p-values for the respective comparisons are shown to the right.

Third, we identified known transcription factor binding motifs enriched within pCRMs using HOMER (Heinz et al., 2010). pCRMs present at the early gastrula stage (stage 10) were significantly enriched (p < 10^-50^ for comparison to random xenlae2 genomic sequence) for binding motifs associated with multiple developmental transcription factors, including Sox, Oct, Zic, Kruppel-like (KLF), Grainyhead-like (GRHL), and Fox family transcription factors (Figure 7D, Supplementary Data 4). Focusing on sites that lose accessibility from stage 10 to stage 12 (ΔpCRMs) identified enrichment of binding motifs only for Sox, Oct, Oct-Sox, and Zic family transcription factors when compared to genomic background (p < 10^-50^); these motifs were also modestly enriched in ΔpCRMs when compared to a background of all pCRMs (Figure 7E, Supplementary Data 4). Sox binding motifs were present in 945 (∼50%) of 1896 surveyed ΔpCRMs, compared with 23% of random genomic sequences and 44% of all pCRMs. Similarly, 427 Oct-Sox motifs were identified in ΔpCRMs (23% of ΔpCRMs vs 6% in random genomic sequence and 17% of pCRMs). The high prevalence of Sox and Oct sites in pCRMs in the early gastrula is consistent with recent ChIP-Seq and DNase-Seq data at the midblastula stage (stage 8.5) of development in the related species *X. tropicalis* (Gentsch et al., 2019). These results provide a basis for further mechanistic investigation into the role of pluripotency factors and the cis-regulatory elements they interact with in the loss of competence.

## Discussion

Competence is a fundamental phenomenon of development and mechanisms underlying loss of competence remain incompletely understood. This study uses ATAC-Seq to define chromatin accessibility over several developmental windows and examines loss of chromatin accessibility as a mechanism for loss of competence. Genome-wide analysis of chromatin accessibility in *Xenopus* demonstrates that promoter accessibility is highly dynamic during gastrula stages. However, the promoters for dorsally localized genes that are direct targets of maternal Wnt signaling, notably *sia1* and *nodal3.1*, are inaccessible at the onset of gastrulation, when competence for the dorsal inductive signal has been lost. This loss of competence is mediated in part by the activity of HDACs, as histone acetylation at the *sia1* and *nodal3.1* promoters is high before the MBT (Blythe et al., 2010) and barely detectable by the late blastula stage, and, furthermore, HDAC inhibition increases acetylation of Wnt target gene promoters and maintains competence to respond to dorsal induction in ventral marginal zone cells, as well as naïve ectoderm. We also determined that mechanisms for loss of competence are context-specific, as competence for mesoderm and neural crest induction are not extended by HDAC inhibition and the promoters for essential regulators of mesoderm (e.g. *tbxt*) and neural crest (*Snai1, Snai2, Twist1*) maintain accessibility after the loss of competence.

Our findings complement recent work investigating the role of chromatin structure in the onset of competence for inductive signals, including evidence that maternal proteins act as pioneer factors to open chromatin at the onset of zygotic transcription (Gentsch et al., 2019; Jacobs et al., 2018; McDaniel et al., 2019) and confer competence for primary germ layer formation in *Xenopus tropicalis* embryos (Gentsch et al., 2019; Paraiso et al., 2019), consistent with the original description of pioneer factors as regulators of developmental competence (Zaret, 1999). With the caveat that we are comparing distinct, though closely related species, and different methods for assessing chromatin accessibility, the similar patterns of accessibility in *X. tropicalis* at the MBT and *X. laevis* gastrulae (Supplementary Figure 1) is intriguing and may suggest that chromatin accessibility is both conserved between these species and is largely maintained through multiple stages of early development. Our findings also complement prior work showing that the earliest steps in dorsal induction include establishment of primed chromatin architecture at dorsal Wnt target genes before ZGA, which in turn allows transcription of epigenetically marked chromatin after ZGA (Blythe et al., 2010). Here we have focused our attention on the loss of competence and ATAC-Seq analysis of chromatin accessibility in *Xenopus*.

Although loss of competence could occur at any level of an inductive signaling pathway, several lines of evidence support that loss of dorsal competence is regulated at the level of Wnt target gene promoters and not through modulation of upstream Wnt signaling (Darken and Wilson, 2001; Hamilton et al., 2001). An inducible activated form of TCF initiates dorsal development when activated before the MBT but not during the late blastula stage (Darken and Wilson, 2001) and an inhibitory form of TCF blocks dorsal development only when induced before the MBT (Blythe et al., 2010; Yang et al., 2002). Furthermore a 0.8 kilobase fragment of the *sia1* promoter follows the same pattern of competence for Wnt signaling as the endogenous *sia1* promoter, indicating that loss of competence is mediated, at least in part, at the level of the promoter (Darken and Wilson, 2001). Evidence that Wnt signaling is intact during and after the loss of competence for dorsal induction includes the observation that ß-catenin can be stabilized by activation of Wnt signaling both before and after the MBT (Darken and Wilson, 2001; Schneider et al., 1996; Schohl and Fagotto, 2002). Furthermore, while dorsal induction in response to maternal Wnt/ß-catenin signaling declines during cleavage and early blastula stages, zygotic Wnt signaling beginning in the blastula stage induces late Wnt target genes to regulate anterior-posterior patterning (Christian and Moon, 1993; Ding et al., 2017; Hamilton et al., 2001; Hoppler et al., 1996; Kjolby and Harland, 2017; Nakamura et al., 2016). Based on this, we conclude that neither the loss of competence for dorsal induction nor the gain of competence for A/P patterning and neural crest induction by Wnt signaling is mediated by changes in the Wnt signal transduction pathway. Instead, our data and previous data support that loss of competence for dorsal induction is mediated at the level of transcription.

Chromatin accessibility is mediated by post-translational modifications of histones, and increased histone acetylation contributes to openness of chromatin and a generally favorable environment for transcription (Shahbazian and Grunstein, 2007). As HDAC inhibition extends the window of competence to induce dorsal development in response to Wnt signaling, we propose that HDACs mediate the loss of competence at dorsal Wnt target genes. The T-cell factor/lymphoid enhancer factor (TCF/LEF) family of transcription factors are downstream mediators of canonical Wnt/ß-catenin signaling that in the absence of Wnt signaling bind diverse corepressors, including Groucho/transducin-like enhancer of split (Gro/TLE) family, C-terminal binding protein (CtBP), Silencing Mediator for Retinoid and Thyroid hormone receptor (SMRT) and Nuclear receptor Co-Repressor (NCoR), and others, all of which recruit HDACs (Cadigan, 2012; Ramakrishnan et al., 2018). In addition, the transcription repressor Kaiso binds to both TCF3 and Wnt-response elements and represses *sia1* expression (Cadigan, 2012; Park et al., 2005; Ramakrishnan et al., 2018). Thus, although the mechanisms of repression by Kaiso are controversial (Cadigan, 2012; Park et al., 2005; Ramakrishnan et al., 2018; Ruzov et al., 2009), the repressive function of Kaiso may also contribute to the loss of competence at dorsal Wnt target genes. We speculate that TCFs bind to Wnt responsive elements throughout competent tissues and could serve two opposing functions. Localized Wnt signaling in future dorsal blastomeres causes ß-catenin accumulation and displacement of corepressors, but in the absence of a Wnt signal, TCF may recruit HDACs that would deaceylate Wnt target gene promoters leading to loss of responsiveness to Wnt signaling. This proposed model is consistent with reports showing that knockdown of maternal *Tcf3* increases *sia1* expression in ventral blastomeres (Houston et al., 2002) and deletion of TCF binding sites in the *sia1* promoter-reporter enhances reporter activity in ventral blastomeres (Brannon et al., 1997), further supporting a repressive function for maternal *Tcf3*. A novel 11 base-pair negative regulatory element (NRE) has recently been identified upstream of multiple Wnt-responsive genes including *sia1* and *nodal3.1* (Kim et al., 2017) and could contribute to loss of competence for dorsal induction.

The regulation of competence for dorsal development may involve additional factors, including alternative TCF proteins with distinct functions after the MBT (Hamilton et al., 2001) and pathway-specific repressors such as BarH like homeobox 2 (Barhl2) (Sena et al., 2019) and Xcad2 (Levy et al., 2002) that appear to function downstream of Wnt-dependent activation of *sia1*. *Barhl2*, which is expressed after the loss of competence to induce dorsal Wnt target genes, stabilizes the interaction of TCF3 (also known as TCF7l1) with Gro/TLE to repress Wnt/ß-catenin target genes in a manner that depends on *Hdac1*. Although *Barhl2* depletion does not affect *sia1* expression, knockdown does expand expression of other organizer genes, suggesting that *Barhl2* acts downstream of *sia1* to limit Wnt-dependent organizer development. Similarly, the homeobox transcription factor *XCad2* also contributes to loss of competence for dorsal induction downstream of *sia1* (Levy et al., 2002). Therefore, *Barhl2* and *Xcad2* may contribute to a mechanism for reinforcing the loss of dorsal competence downstream of the initial response to Wnt/ß-catenin signaling at *sia1*.

Although HDAC inhibition extends competence for dorsal development, it is not sufficient to extend competence for mesoderm induction in ectodermal explants, consistent with a report that TSA did not expand expression of mesendodermal genes in intact embryos (Gao et al., 2016). Alternatively, loss of competence for mesoderm induction could be mediated by tissue-specific transcription repressors. For example, overexpression of *Ascl1*, a neural regulator, suppresses *tbxt* and mesendodermal genes in an HDAC-dependent manner (Gao et al., 2016). Other tissue-specific suppressors of mesodermal gene expression may also mediate loss of mesodermal competence, including *pou5f3.1* (*oct91/pou91*) (Henig et al., 1998), *pou5f3.2 (oct-25*), and *pou5f3.3* (*oct-60*), which inhibit mesoderm induction by both FGF and activin/nodal family members (Cao et al., 2006).

ATAC peaks were not reduced at multiple mesodermal gene promoters after the loss of mesodermal competence, including *tbxt*, a direct target of FGF signaling and a master regulator of mesoderm induction. Although histone deacetylation at other genomic regions could mediate loss of competence, global inhibition of HDACs did not extend competence for FGF. Nevertheless, HDAC-independent changes in chromatin architecture could mediate loss of competence to induce mesoderm. For example, Steinbach and colleagues showed that accumulation of histone H1 after the MBT leads to loss of competence for mesoderm induction (Steinbach et al., 1998). Histone H1 accumulation causes chromatin compaction (Happel and Doenecke, 2009) and correlates with deposition of the repressive mark H3K9me3 (Cao et al., 2013). Thus, zygotic expression of H1 could represent an HDAC-independent mechanism to reduce chromatin accessibility and competence for mesoderm inducing signals. In addition, Jarid2/Jumonji, a component and antagonist of the Polycomb Repressive Complex 2 (PRC2), is required for mesodermal gene expression (Peng et al., 2009). Knockdown of *Jarid2* in *Xenopus* impairs expression of *tbxt* and blocks mesoderm induction by activin, suggesting that increased PRC2 activity suppresses mesodermal gene expression and that modulation of Jarid2 function could contribute to the loss of competence during the gastrula stage. PRC2 deposits the repressive modification H3K27me3, which has also been proposed to contribute to loss of competence to mesoderm induction by Smad2 in zebrafish (Shiomi et al., 2017).

Similarly, HDAC-*independent* modulation of chromatin architecture could mediate the loss of competence for neural crest induction. The histone methyltransferase Prdm12 methylates H3K9 at the *snai2* and *foxd3* promoters and *prdm12* overexpression suppresses neural crest induction (Matsukawa et al., 2015). PRC2 also interacts with Snail2/Slug, methylates H3K27 at Snail2/Slug target gene promoters, and is required for neural crest specification and migration downstream of *snail1* and *snail2* (Tien et al., 2015).

Although HDAC inhibition did not extend competence for mesoderm or neural crest induction, global inhibition of HDACs at early stages may interfere with later developmental events and confound our analysis of inductive competence. For example, we observed that prior exposure to TSA impaired neural crest induction by Wnt activation at stages that are competent to respond, consistent with a report showing that HDAC activity is necessary for Wnt-mediated neural crest induction (Rao and LaBonne, 2018). Additionally, TSA treatment in ectodermal explants (at concentrations 5-fold higher than used here) decreases expression of pluripotency markers and significantly increases expression of lineage-specific markers, including the mesodermal genes *tbxt* and *myod1*, as well as endodermal and neural specific genes (Rao and LaBonne, 2018). HDAC inhibition at early developmental stages may therefore extend competence windows so that naïve cells respond inappropriately to endogenous signals, diverting them into inappropriate lineage-restricted states and blocking induction by later signals, such as mesoderm or neural crest inducing signals.

Thus, changes in accessibility that we observe during gastrulation may still contribute to loss of competence to respond to developmental signals at these promoters and at promoter-distal cis-regulatory sites. It is intriguing, therefore, that binding sites for Oct and Sox family pluripotency factors are highly enriched in putative cis-regulatory modules during gastrulation, including pCRMs that lose accessibility during gastrulation, as these factors were recently shown to open chromatin and play a crucial role in establishing competence for inductive signals at the earlier stage of zygotic gene activation in *X. tropicalis* (Gentsch et al., 2019). However, whether closing of regulatory sites during loss of competence is mediated by changes in expression, activity, or accessibility of pluripotency factors remains a subject for future study.

## Conclusions

In summary, we employed ATAC-Seq to identify changes in chromatin accessibility associated with competence to respond to inductive signals in the early embryo. We find that competence to respond to the earliest inductive signal, Wnt pathway-dependent induction of the Spemann organizer, is regulated by changes in chromatin architecture, with the loss of competence mediated, at least in part, by deacetylation of Wnt target gene promoters. We also find that, while the mechanisms for regulating competence appear to be context-dependent, binding sites for pluripotency factors are enriched in putative regulatory elements at the gastrula stage including sites that lose accessibility in parallel with tissue specification and loss of competence during the gastrula stage. Future work will address the mechanisms that regulate chromatin accessibility and the impact of changes in chromatin structure on developmental competence.

## Supporting information

description of supplemental files

## Acknowledgements

We greatly appreciate Zhijun Duan for providing technical guidance with ATAC-seq library construction. We thank We thank Parisha Shah, Gert Veenstra, Douglas Epstein, Daniel Kessler, Mary Mullins, Montserrat Anguera, Melinda Snitow, and David Klein for helpful advice and Linyang Ju for advice and generously supplying Tn5 transposase. M.E. was supported in part by the Medical Scientist Training Program at the Perelman School of Medicine at the University of Pennsylvania. This work was supported by grants from the National Institutes of Health including 1R01GM115517 and 1R01HL141759 (PSK); R01GM111816 and R35GM131810 (JY); R01GM132438 (KZ); T32GM007170, F31GM116588, and T32HL007439 (ME).

---

Supplemental data available at:

https://data.mendeley.com/datasets/fbkvhzbyy5/draft?a=fb936847-4075-4d6e-93fa-f8e2e412378e

1. Data description ATAC-seq in Xenopus (Esmaeili et al), DOI: http://dx.doi.org/10.17632/fbkvhzbyy5.2#file-2736ae60-b4dd-42b2-9c46-06345579b77a
2. ATAC-seq analysis for Xenopus stage 10 and stage 12, DOI: http://dx.doi.org/10.17632/fbkvhzbyy5.2#file-e749e6b6-0436-4ecb-9eed-cfdc95672b77
3. ATAC-seq vs expression Xenopus stage 10.xlsx, DOI: http://dx.doi.org/10.17632/fbkvhzbyy5.2#file-833324f4-4b89-4090-a1dc-6678f6d4ba75
4. Reduced accessibility at putative cis-regulatory modules in Xenopus gastrula stage ectoderm, DOI: http://dx.doi.org/10.17632/fbkvhzbyy5.2#file-13c5f4bf-c61e-4d48-8c7a-c35c4796b0c6
5. Motif analysis for putative cis-regulatory modules in Xenopus gastrula, DOI: http://dx.doi.org/10.17632/fbkvhzbyy5.2#file-97152196-5767-402e-a6e2-2c9e0ad9079e
6. Comparison of ATAC-Seq in X. laevis stage 10 to Dnase-Seq in X. tropicalis stage 8.5

Supplementary data for Esmaeili et al, “Chromatin accessibility and histone acetylation in the regulation of competence in early development”

1. Supplementary Data 1: ATAC-seq analysis for Xenopus stage 10 and stage 12

a. ATAC-seq analysis stage 10 and 12 with p300 overlap.
b. Genome-wide distribution of ATAC-seq peaks.
c. List of promoters for which accessibility changes ≥ 2-fold.
d. Putative cis-regulatory modules (pCRMs).
e. ΔpCRMs: pCRMs with reduced accessibility from stage 10 to stage 12.
2. Supplementary Data 2: ATAC-seq vs expression Xenopus stage 10

a. Comparison of accessibility at promoters annotated in xenlae2 with expression at stage 10 by microarray (Livigni et al 2013).
b. List of genes based on high vs low promoter accessibility and high vs low expression at stage 10.
c. DAVID functional gene annotation.
3. Supplementary Data 3: Reduced accessibility at putative cis-regulatory modules in Xenopus gastrula stage ectoderm

a. DAVID functional gene annotation for ΔpCRMs.
b. List of gene names for nearest TSS within 100kb of ΔpCRM.
4. Supplementary Data 4: Motif analysis for putative cis-regulatory modules in Xenopus gastrula

a. pCRMs compared to genome background.
b. ΔpCRMs compared to genome background.
c. ΔpCRMs compared to pCRM background.
5. Supplementary Figure 1: Comparison of ATAC-Seq gastrula stage (laevis) and Dnase-Seq MBT (tropicalis)

a. Tracks for selected genes from X. laevis ATAC-Seq at stage 10 gastrula (Esmaeili et al) to X. tropicalis Dnase-Seq at stage 8.5 (MBT) from Gentsch et al 2019.

**Supplementary Figure 1:**
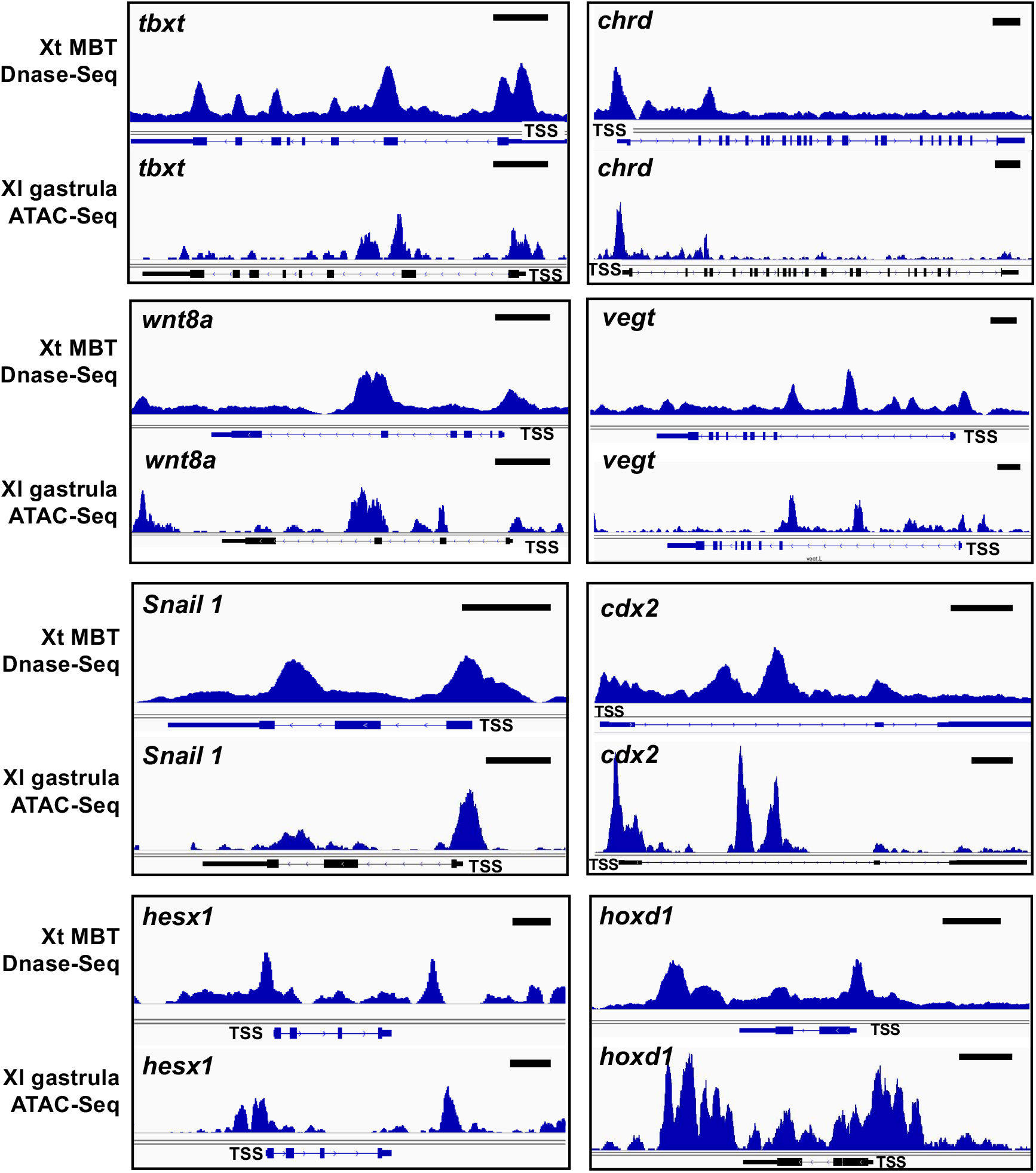
Chromatin accessibility by ATAC-Seq (*X. laevis* gastrula) vs Dnase-Seq (*X. tropicalis*, MBT) ATAC-Seq tracks from stage 10 (early gastrula) *Xenopus laevis* ectodermal explants (GSE GSE76059) were compared to Dnase-Seq tracks from stage 8.5 (MBT) *Xenopus tropicalis* embryos from (Gentsch et al 2019; GSE113186). Tracks were viewed in IGV. Scale bars indicate 1kb. Y-axes were adjusted for each gene to allow side-by-side comparisons and do not reflect relative intensity of signals between different genes, species, stage, or method of analysis. TSS: transcription start site. “Xl”: *Xenopus laevis*; “Xt: *Xenopus topicalis*.

## KEY RESOURCES TABLE

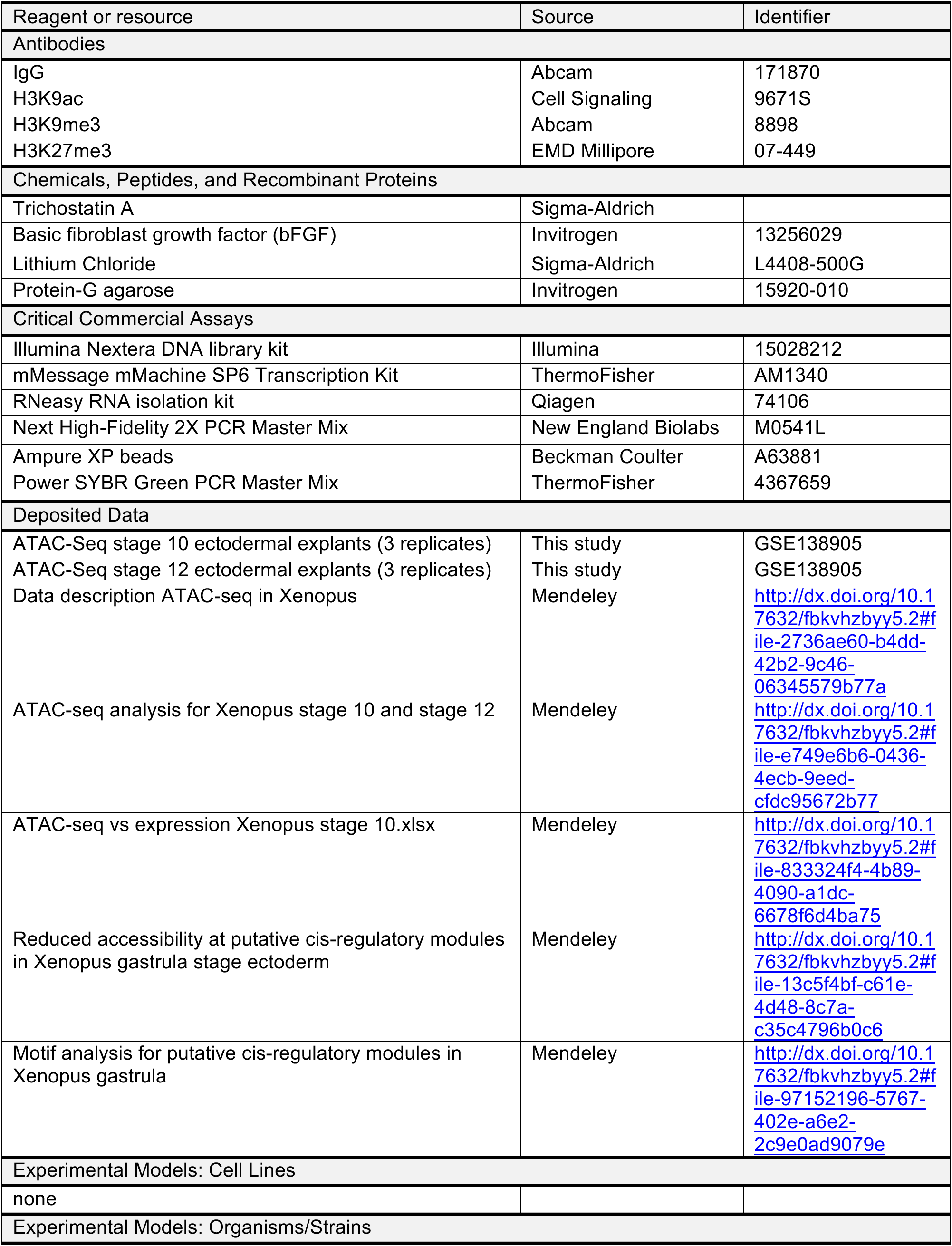
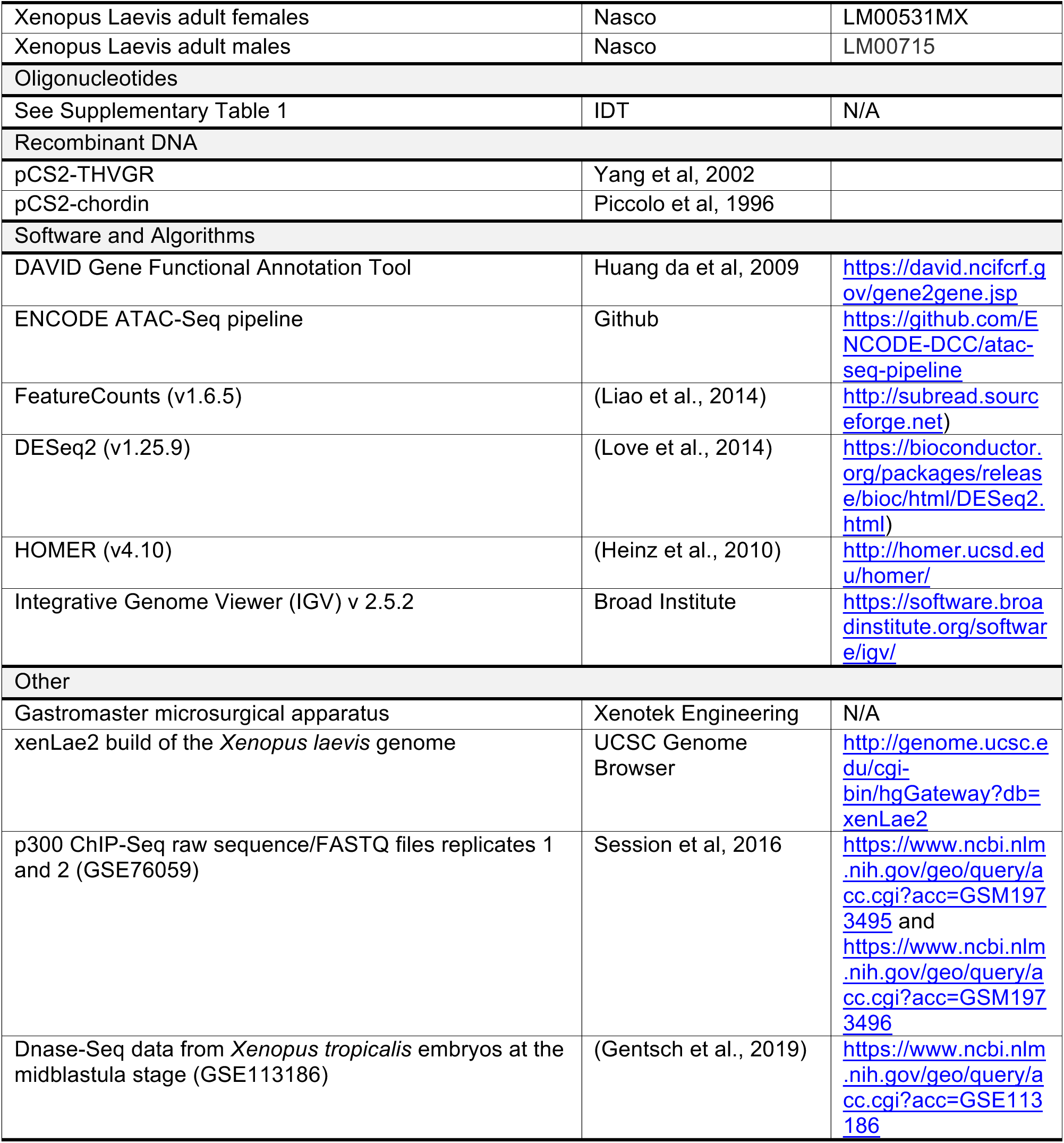

## Notes

#### Summary of Updates

Corrected error in Figure 7 (duplicated panels).

https://data.mendeley.com/datasets/fbkvhzbyy5/1

## References

Akkers, R.C., van Heeringen, S.J., Jacobi, U.G., Janssen-Megens, E.M., Francoijs, K.J., Stunnenberg, H.G., Veenstra, G.J., 2009. A hierarchy of H3K4me3 and H3K27me3 acquisition in spatial gene regulation in Xenopus embryos. Dev Cell 17, 425–434.

Asashima, M., Nakano, H., Shimada, K., Kinoshita, K., Ishii, K., Shibai, H., Ueno, N., 1990. Mesodermal induction in early amphibian embryos by activin A (erythroid differentiation factor). Roux’s Arch. Dev. Biol. 198, 330–335.

Aybar, M.J., Mayor, R., 2002. Early induction of neural crest cells: lessons learned from frog, fish and chick. Current Opinion in Genetics & Development 12, 452–458.

Bae, S., Reid, C.D., Kessler, D.S., 2011. Siamois and Twin are redundant and essential in formation of the Spemann organizer. Developmental biology 352, 367–381.

Beisel, C., Paro, R., 2011. Silencing chromatin: comparing modes and mechanisms. Nat Rev Genet 12, 123–135.

Blythe, S.A., Cha, S.W., Tadjuidje, E., Heasman, J., Klein, P.S., 2010. beta-Catenin primes organizer gene expression by recruiting a histone H3 arginine 8 methyltransferase, Prmt2. Developmental Cell 19, 220–231.

Blythe, S.A., Reid, C.D., Kessler, D.S., Klein, P.S., 2009. Chromatin immunoprecipitation in early Xenopus laevis embryos. Developmental Dynamics 238, 1422–1432.

Brannon, M., Gomperts, M., Sumoy, L., Moon, R.T., Kimelman, D., 1997. A beta-catenin/XTcf-3 complex binds to the siamois promoter to regulate dorsal axis specification in Xenopus. Genes & development 11, 2359–2370.

Bright, A.R., Veenstra, G.J.C., 2019. Assay for Transposase-Accessible Chromatin-Sequencing Using Xenopus Embryos. Cold Spring Harb Protoc 2019, pdb prot098327.

Buenrostro, J.D., Giresi, P.G., Zaba, L.C., Chang, H.Y., Greenleaf, W.J., 2013. Transposition of native chromatin for fast and sensitive epigenomic profiling of open chromatin, DNA-binding proteins and nucleosome position. Nat Methods 10, 1213–1218.

Cadigan, K.M., 2012. TCFs and Wnt/beta-catenin signaling: more than one way to throw the switch. Curr Top Dev Biol 98, 1–34.

Cao, K., Lailler, N., Zhang, Y., Kumar, A., Uppal, K., Liu, Z., Lee, E.K., Wu, H., Medrzycki, M., Pan, C., Ho, P.Y., Cooper, G.P., Jr., Dong, X., Bock, C., Bouhassira, E.E., Fan, Y., 2013. High-resolution mapping of h1 linker histone variants in embryonic stem cells. PLoS Genet 9, e1003417.

Cao, Y., Siegel, D., Knochel, W., 2006. Xenopus POU factors of subclass V inhibit activin/nodal signaling during gastrulation. Mech Dev 123, 614–625.

Carnac, G., Kodjabachian, L., Gurdon, J.B., Lemaire, P., 1996. The homeobox gene Siamois is a target of the Wnt dorsalisation pathway and triggers organiser activity in the absence of mesoderm. Development 122, 3055–3065.

Cha, S.W., Tadjuidje, E., Tao, Q., Wylie, C., Heasman, J., 2008. Wnt5a and Wnt11 interact in a maternal Dkk1-regulated fashion to activate both canonical and non-canonical signaling in Xenopus axis formation. Development 135, 3719–3729.

Charney, R.M., Forouzmand, E., Cho, J.S., Cheung, J., Paraiso, K.D., Yasuoka, Y., Takahashi, S., Taira, M., Blitz, I.L., Xie, X., Cho, K.W., 2017. Foxh1 Occupies cis-Regulatory Modules Prior to Dynamic Transcription Factor Interactions Controlling the Mesendoderm Gene Program. Dev Cell 40, 595–607 e594.

Christian, J.L., Moon, R.T., 1993. Interactions between Xwnt-8 and Spemann organizer signaling pathways generate dorsoventral pattern in the embryonic mesoderm of Xenopus. Genes & development 7, 13–28.

Dale, L., Smith, J.C., Slack, J.M.W., 1985. Mesoderm induction in *Xenopus laevis:* a quantitative study using a cell lineage label and tissue specific antibodies. J.Embryol.exp.Morph. 89, 289–312.

Darken, R.S., Wilson, P.A., 2001. Axis induction by wnt signaling: Target promoter responsiveness regulates competence. Developmental biology 234, 42–54.

De Robertis, E.M., Larrain, J., Oelgeschlager, M., Wessely, O., 2000. The establishment of Spemann’s organizer and patterning of the vertebrate embryo. Nature Reviews Genetics 1, 171–181.

Ding, Y., Ploper, D., Sosa, E.A., Colozza, G., Moriyama, Y., Benitez, M.D., Zhang, K., Merkurjev, D., De Robertis, E.M., 2017. Spemann organizer transcriptome induction by early beta-catenin, Wnt, Nodal, and Siamois signals in Xenopus laevis. Proceedings of the National Academy of Sciences of the United States of America 114, E3081–E3090.

Elurbe, D.M., Paranjpe, S.S., Georgiou, G., van Kruijsbergen, I., Bogdanovic, O., Gibeaux, R., Heald, R., Lister, R., Huynen, M.A., van Heeringen, S.J., Veenstra, G.J.C., 2017. Regulatory remodeling in the allo-tetraploid frog Xenopus laevis. Genome Biol 18, 198.

Fagotto, F., Guger, K., Gumbiner, B.M., 1997. Induction of the primary dorsalizing center in Xenopus by the Wnt/GSK/beta-catenin signaling pathway, but not by Vg1, Activin or Noggin. Development 124, 453–460.

Fredieu, J.R., Cui, Y., Maier, D., Danilchik, M.V., Christian, J.L., 1997. Xwnt-8 and lithium can act upon either dorsal mesodermal or neurectodermal cells to cause a loss of forebrain in Xenopus embryos. Developmental biology 186, 100–114.

Gao, L., Zhu, X., Chen, G., Ma, X., Zhang, Y., Khand, A.A., Shi, H., Gu, F., Lin, H., Chen, Y., Zhang, H., He, L., Tao, Q., 2016. A novel role for Ascl1 in the regulation of mesendoderm formation via HDAC-dependent antagonism of VegT. Development 143, 492–503.

Gentsch, G.E., Spruce, T., Owens, N.D.L., Smith, J.C., 2019. Maternal pluripotency factors initiate extensive chromatin remodelling to predefine first response to inductive signals. Nat Commun 10, 4269.

Green, J.B., Howes, G., Symes, K., Cooke, J., Smith, J.C., 1990. The biological effects of XTC-MIF: quantitative comparison with *Xenopus* bFGF. Development 108, 173–183.

Gupta, R., Wills, A., Ucar, D., Baker, J., 2014. Developmental enhancers are marked independently of zygotic Nodal signals in Xenopus. Developmental biology 395, 38–49.

Gurdon, J.B., Fairman, S., Mohun, T.J., Brennan, S., 1985. Activation of muscle specific actin genes in *Xenopus* development by an induction between animal and vegetal cells of a blastula. Cell 41, 913–922.

Hamburger, V., 1988. The Heritage of Experimental Embryology: Hans Spemann and the organizer. Oxford University Press, New York.

Hamilton, F.S., Wheeler, G.N., Hoppler, S., 2001. Difference in XTcf-3 dependency accounts for change in response to beta-catenin-mediated Wnt signalling in Xenopus blastula. Development 128, 2063–2073.

Happel, N., Doenecke, D., 2009. Histone H1 and its isoforms: contribution to chromatin structure and function. Gene 431, 1–12.

Heasman, J., Kofron, M., Wylie, C., 2000. Beta-catenin signaling activity dissected in the early Xenopus embryo: a novel antisense approach. Developmental biology 222, 124–134.

Hecht, A., Vleminckx, K., Stemmler, M.P., van Roy, F., Kemler, R., 2000. The p300/CBP acetyltransferases function as transcriptional coactivators of beta-catenin in vertebrates. Embo J 19, 1839–1850.

Heinz, S., Benner, C., Spann, N., Bertolino, E., Lin, Y.C., Laslo, P., Cheng, J.X., Murre, C., Singh, H., Glass, C.K., 2010. Simple combinations of lineage-determining transcription factors prime cis-regulatory elements required for macrophage and B cell identities. Molecular cell 38, 576–589.

Henig, C., Elias, S., Frank, D., 1998. A POU protein regulates mesodermal competence to FGF in Xenopus. Mech Dev 71, 131–142.

Herberg, S., Simeone, A., Oikawa, M., Jullien, J., Bradshaw, C.R., Teperek, M., Gurdon, J., Miyamoto, K., 2015. Histone H3 lysine 9 trimethylation is required for suppressing the expression of an embryonically activated retrotransposon in Xenopus laevis. Sci Rep 5, 14236.

Hontelez, S., van Kruijsbergen, I., Georgiou, G., van Heeringen, S.J., Bogdanovic, O., Lister, R., Veenstra, G.J.C., 2015. Embryonic transcription is controlled by maternally defined chromatin state. Nat Commun 6, 10148.

Hoppler, S., Brown, J.D., Moon, R.T., 1996. Expression of a dominant-negative Wnt blocks induction of MyoD in Xenopus embryos. Genes & development 10, 2805–2817.

Houston, D.W., Kofron, M., Resnik, E., Langland, R., Destree, O., Wylie, C., Heasman, J., 2002. Repression of organizer genes in dorsal and ventral Xenopus cells mediated by maternal XTcf3. Development 129, 4015–4025.

Huang da, W., Sherman, B.T., Lempicki, R.A., 2009. Systematic and integrative analysis of large gene lists using DAVID bioinformatics resources. Nat Protoc 4, 44–57.

Ishibashi, H., Matsumura, N., Hanafusa, H., Matsumoto, K., De Robertis, E.M., Kuroda, H., 2008. Expression of Siamois and Twin in the blastula Chordin/Noggin signaling center is required for brain formation in Xenopus laevis embryos. Mech Dev 125, 58–66.

Jacobs, J., Atkins, M., Davie, K., Imrichova, H., Romanelli, L., Christiaens, V., Hulselmans, G., Potier, D., Wouters, J., Taskiran, II, Paciello, G., Gonzalez-Blas, C.B., Koldere, D., Aibar, S., Halder, G., Aerts, S., 2018. The transcription factor Grainy head primes epithelial enhancers for spatiotemporal activation by displacing nucleosomes. Nature genetics 50, 1011–1020.

Jones, E.A., Woodland, H.R., 1987. The development of animal cap cells in Xenopus: a measure of the start of animal cap competence to form mesoderm. Development 101, 557–563.

Kao, K.R., Elinson, R.P., 1988. The entire mesodermal mantle behaves as Spemann’s organizer in dorsoanterior enhanced *Xenopus laevis* embryos. Developmental biology 127, 64–77.

Kao, K.R., Masui, Y., Elinson, R.P., 1986. Lithium-induced respecification of pattern in Xenopus laevis embryos. Nature 322, 371–373.

Karimi, K., Fortriede, J.D., Lotay, V.S., Burns, K.A., Wang, D.Z., Fisher, M.E., Pells, T.J., James-Zorn, C., Wang, Y., Ponferrada, V.G., Chu, S., Chaturvedi, P., Zorn, A.M., Vize, P.D., 2018. Xenbase: a genomic, epigenomic and transcriptomic model organism database. Nucleic Acids Res 46, D861–D868.

Kengaku, M., Okamoto, H., 1993. Basic fibroblast growth factor induces differentiation of neural tube and neural crest lineages of cultured ectoderm cells from Xenopus gastrula. Development 119, 1067–1078.

Kessler, D.S., 1997. Siamois is required for formation of Spemann’s organizer. Proceedings of the National Academy of Sciences of the United States of America 94, 13017–13022.

Kim, K., Cho, J., Hilzinger, T.S., Nunns, H., Liu, A., Ryba, B.E., Goentoro, L., 2017. Two-Element Transcriptional Regulation in the Canonical Wnt Pathway. Curr Biol 27, 2357–2364 e2355.

Kjolby, R.A.S., Harland, R.M., 2017. Genome-wide identification of Wnt/beta-catenin transcriptional targets during Xenopus gastrulation. Developmental biology 426, 165–175.

Kodjabachian, L., Lemaire, P., 2001. Siamois functions in the early blastula to induce Spemann’s organiser. Mech Dev 108, 71–79.

Lamb, T.M., Knecht, A.K., Smith, W.C., Stachel, S.E., Economides, A.N., Stahl, N., Yancopolous, G.D., Harland, R.M., 1993. Neural Induction by the Secreted Polypeptide Noggin. Science 262, 713–718.

Laurent, M.N., Blitz, I.L., Hashimoto, C., Rothbacher, U., Cho, K.W., 1997. The Xenopus homeobox gene twin mediates Wnt induction of goosecoid in establishment of Spemann’s organizer. Development 124, 4905–4916.

Lee, M.T., Bonneau, A.R., Takacs, C.M., Bazzini, A.A., DiVito, K.R., Fleming, E.S., Giraldez, A.J., 2013. Nanog, Pou5f1 and SoxB1 activate zygotic gene expression during the maternal-to-zygotic transition. Nature 503, 360–364.

Leichsenring, M., Maes, J., Mossner, R., Driever, W., Onichtchouk, D., 2013. Pou5f1 transcription factor controls zygotic gene activation in vertebrates. Science 341, 1005–1009.

Lemaire, P., Garrett, N., Gurdon, J.B., 1995. Expression cloning of Siamois, a Xenopus homeobox gene expressed in dorsal-vegetal cells of blastulae and able to induce a complete secondary axis. Cell 81, 85–94.

Levy, V., Marom, K., Zins, S., Koutsia, N., Yelin, R., Fainsod, A., 2002. The competence of marginal zone cells to become Spemann’s organizer is controlled by Xcad2. Developmental biology 248, 40–51.

Liao, Y., Smyth, G.K., Shi, W., 2014. featureCounts: an efficient general purpose program for assigning sequence reads to genomic features. Bioinformatics 30, 923–930.

Livigni, A., Peradziryi, H., Sharov, A.A., Chia, G., Hammachi, F., Migueles, R.P., Sukparangsi, W., Pernagallo, S., Bradley, M., Nichols, J., Ko, M.S.H., Brickman, J.M., 2013. A conserved Oct4/POUV-dependent network links adhesion and migration to progenitor maintenance. Curr Biol 23, 2233–2244.

Love, M.I., Huber, W., Anders, S., 2014. Moderated estimation of fold change and dispersion for RNA-seq data with DESeq2. Genome Biol 15, 550.

Lu, F.I., Thisse, C., Thisse, B., 2011. Identification and mechanism of regulation of the zebrafish dorsal determinant. Proceedings of the National Academy of Sciences of the United States of America 108, 15876–15880.

Mancilla, A., Mayor, R., 1996. Neural crest formation in Xenopus laevis: mechanisms of Xslug induction. Developmental biology 177, 580–589.

Matsukawa, S., Miwata, K., Asashima, M., Michiue, T., 2015. The requirement of histone modification by PRDM12 and Kdm4a for the development of pre-placodal ectoderm and neural crest in Xenopus. Developmental biology 399, 164–176.

McDaniel, S.L., Gibson, T.J., Schulz, K.N., Fernandez Garcia, M., Nevil, M., Jain, S.U., Lewis, P.W., Zaret, K.S., Harrison, M.M., 2019. Continued Activity of the Pioneer Factor Zelda Is Required to Drive Zygotic Genome Activation. Molecular cell 74, 185–195 e184.

Moon, R.T., Kimelman, D., 1998. From cortical rotation to organizer gene expression: toward a molecular explanation of axis specification in Xenopus. Bioessays 20, 536–545.

Nakamura, O., Hayashi, Y., Asashima, M., 1978. A half century from Spemann - Historical review of studies on the organizer, in: Toivonen, O.N.S. (Ed.), Organizer - A milestone of a half-century from Spemann. North Holland Biochemical Press, Elsevier, pp. 1–47.

Nakamura, O., Takasaki, H., Ishihara, M., 1970. Formation of the organizer from combinations of presumptive ectoderm and endoderm. I. Proc. Japan Acad. 47, 313–331.

Nakamura, Y., de Paiva Alves, E., Veenstra, G.J., Hoppler, S., 2016. Tissue- and stage-specific Wnt target gene expression is controlled subsequent to beta-catenin recruitment to cis-regulatory modules. Development 143, 1914–1925.

Nieuwkoop, P.D., Faber, J., 1967. Normal Table of Xenopus laevis (Daudin), Second edition ed. North Holland Publishing Company, Amsterdam.

Paraiso, K.D., Blitz, I.L., Coley, M., Cheung, J., Sudou, N., Taira, M., Cho, K.W.Y., 2019. Endodermal Maternal Transcription Factors Establish Super-Enhancers during Zygotic Genome Activation. Cell Rep 27, 2962–2977 e2965.

Park, J.I., Kim, S.W., Lyons, J.P., Ji, H., Nguyen, T.T., Cho, K., Barton, M.C., Deroo, T., Vleminckx, K., Moon, R.T., McCrea, P.D., 2005. Kaiso/p120-catenin and TCF/beta-catenin complexes coordinately regulate canonical Wnt gene targets, Developmental Cell, pp. 843–854.

Peng, J.C., Valouev, A., Swigut, T., Zhang, J., Zhao, Y., Sidow, A., Wysocka, J., 2009. Jarid2/Jumonji coordinates control of PRC2 enzymatic activity and target gene occupancy in pluripotent cells. Cell 139, 1290–1302.

Perino, M., Veenstra, G.J., 2016. Chromatin Control of Developmental Dynamics and Plasticity. Dev Cell 38, 610–620.

Piccolo, S., Sasai, Y., Lu, B., Derobertis, E.M., 1996. Dorsoventral Patterning In Xenopus: Inhibition Of Ventral Signals By Direct Binding Of Chordin to Bmp-4. Cell 0086, 589–598.

Ramakrishnan, A.B., Sinha, A., Fan, V.B., Cadigan, K.M., 2018. The Wnt Transcriptional Switch: TLE Removal or Inactivation? Bioessays 40, 1–6.

Rao, A., LaBonne, C., 2018. Histone deacetylase activity has an essential role in establishing and maintaining the vertebrate neural crest. Development 145, 1–11.

Robinson, J.T., Thorvaldsdottir, H., Winckler, W., Guttman, M., Lander, E.S., Getz, G., Mesirov, J.P., 2011. Integrative genomics viewer. Nature biotechnology 29, 24–26.

Rosa, F., Roberts, A.B., Danielpour, D., Dart, L.L., Sporn, M.B., Dawid, I.B., 1988. Mesoderm induction in amphibians: the role of TGF-b2 like factors. Science 239, 783–785.

Ruzov, A., Hackett, J.A., Prokhortchouk, A., Reddington, J.P., Madej, M.J., Dunican, D.S., Prokhortchouk, E., Pennings, S., Meehan, R.R., 2009. The interaction of xKaiso with xTcf3: a revised model for integration of epigenetic and Wnt signalling pathways. Development 136, 723–727.

Saint-Jeannet, J.P., He, X., Varmus, H.E., Dawid, I.B., 1997. Regulation of dorsal fate in the neuraxis by Wnt-1 and Wnt-3a. Proceedings of the National Academy of Sciences of the United States of America 94, 13713–13718.

Schneider, S., Steinbeisser, H., Warga, R.M., Hausen, P., 1996. Beta-catenin translocation into nuclei demarcates the dorsalizing centers in frog and fish embryos. Mech Dev 57, 191–198.

Schneider, T.D., Arteaga-Salas, J.M., Mentele, E., David, R., Nicetto, D., Imhof, A., Rupp, R.A., 2011. Stage-specific histone modification profiles reveal global transitions in the Xenopus embryonic epigenome. PloS one 6, e22548.

Schohl, A., Fagotto, F., 2002. Beta-catenin, MAPK and Smad signaling during early Xenopus development. Development 129, 37–52.

Sena, E., Rocques, N., Borday, C., Muhamad Amin, H.S., Parain, K., Sitbon, D., Chesneau, A., Durand, B.C., 2019. Barhl2 maintains T cell factors as repressors and thereby switches off the Wnt/beta-Catenin response driving Spemann organizer formation. Development 146, 1–16.

Session, A.M., Uno, Y., Kwon, T., Chapman, J.A., Toyoda, A., Takahashi, S., Fukui, A., Hikosaka, A., Suzuki, A., Kondo, M., van Heeringen, S.J., Quigley, I., Heinz, S., Ogino, H., Ochi, H., Hellsten, U., Lyons, J.B., Simakov, O., Putnam, N., Stites, J., Kuroki, Y., Tanaka, T., Michiue, T., Watanabe, M., Bogdanovic, O., Lister, R., Georgiou, G., Paranjpe, S.S., van Kruijsbergen, I., Shu, S., Carlson, J., Kinoshita, T., Ohta, Y., Mawaribuchi, S., Jenkins, J., Grimwood, J., Schmutz, J., Mitros, T., Mozaffari, S.V., Suzuki, Y., Haramoto, Y., Yamamoto, T.S., Takagi, C., Heald, R., Miller, K., Haudenschild, C., Kitzman, J., Nakayama, T., Izutsu, Y., Robert, J., Fortriede, J., Burns, K., Lotay, V., Karimi, K., Yasuoka, Y., Dichmann, D.S., Flajnik, M.F., Houston, D.W., Shendure, J., DuPasquier, L., Vize, P.D., Zorn, A.M., Ito, M., Marcotte, E.M., Wallingford, J.B., Ito, Y., Asashima, M., Ueno, N., Matsuda, Y., Veenstra, G.J., Fujiyama, A., Harland, R.M., Taira, M., Rokhsar, D.S., 2016. Genome evolution in the allotetraploid frog Xenopus laevis. Nature 538, 336–343.

Shahbazian, M.D., Grunstein, M., 2007. Functions of site-specific histone acetylation and deacetylation. Annu Rev Biochem 76, 75–100.

Shiomi, T., Muto, A., Hozumi, S., Kimura, H., Kikuchi, Y., 2017. Histone H3 Lysine 27 Trimethylation Leads to Loss of Mesendodermal Competence During Gastrulation in Zebrafish Ectodermal Cells. Zoolog Sci 34, 64–71.

Sive, H., Grainger, R., Harland, R., 2000. Early Development of Xenopus laevis; A Laboratory Manual, 1st ed. Cold Spring Harbor Press, Cold Spring Harbor.

Skirkanich, J., Luxardi, G., Yang, J., Kodjabachian, L., Klein, P.S., 2011. An essential role for transcription before the MBT in Xenopus laevis. Developmental biology 357, 478–491.

Slack, J.M.W., Darlington, B.G., Heath, J.K., Godsave, S.F., 1987. Mesoderm induction in early *Xenopus* embryos by heparin-binding growth factors. Nature 326, 197–200.

Smith, W.C., McKendry, R., Ribisi, S., Jr., Harland, R.M., 1995. A nodal-related gene defines a physical and functional domain within the Spemann organizer. Cell 82, 37–46.

Sokol, S., Wong, G.G., Melton, D.A., 1990. A mouse macrophage factor induces head structures and organizes a body axis in *Xenopus*. Science 249, 561–564.

Sokol, S.Y., 1999. Wnt signaling and dorso-ventral axis specification in vertebrates. Curr Opin Genet Dev 9, 405–410.

Spemann, H., 1938. Embryonic Development and Induction. Yale University Press, New York.

Starks, R.R., Biswas, A., Jain, A., Tuteja, G., 2019. Combined analysis of dissimilar promoter accessibility and gene expression profiles identifies tissue-specific genes and actively repressed networks. Epigenetics Chromatin 12, 16.

Steinbach, O.C., Ulshofer, A., Authaler, A., Rupp, R.A., 1998. Temporal restriction of MyoD induction and autocatalysis during Xenopus mesoderm formation. Developmental biology 202, 280–292.

Sudarwati, S., Nieuwkoop, P.D., 1971. Mesoderm formation in the anuran Xenopus laevis (Daudin). Wilhelm Roux Archi. EntwMech. Org. 166, 189–204.

Tao, Q., Yokota, C., Puck, H., Kofron, M., Birsoy, B., Yan, D., Asashima, M., Wylie, C.C., Lin, X., Heasman, J., 2005. Maternal wnt11 activates the canonical wnt signaling pathway required for axis formation in Xenopus embryos. Cell 120, 857–871.

Tien, C.L., Jones, A., Wang, H., Gerigk, M., Nozell, S., Chang, C., 2015. Snail2/Slug cooperates with Polycomb repressive complex 2 (PRC2) to regulate neural crest development. Development 142, 722–731.

Waddington, C.H., 1940. Competence, Organisers and Genes. Cambridge University Press, London, pp. 41–55.

Wu, J., Saint-Jeannet, J.P., Klein, P.S., 2003. Wnt-frizzled signaling in neural crest formation. Trends in Neurosciences 26, 40–45.

Wu, J., Yang, J., Klein, P.S., 2005. Neural crest induction by the canonical Wnt pathway can be dissociated from anterior-posterior neural patterning in Xenopus. Developmental biology 279, 220–232.

Yamaguchi, Y., Shinagawa, A., 1989. Marked alteration at midblastula transition in the effect of lithium on formation of the larval body …. Dev Growth Differ 31, 531–541.

Yan, L., Chen, J., Zhu, X., Sun, J., Wu, X., Shen, W., Zhang, W., Tao, Q., Meng, A., 2018. Maternal Huluwa dictates the embryonic body axis through beta-catenin in vertebrates. Science 362, 1–11.

Yang, J., Tan, C., Darken, R.S., Wilson, P.A., Klein, P.S., 2002. Beta-catenin/Tcf-regulated transcription prior to the midblastula transition. Development 129, 5743–5752.

Yang-Snyder, J., Miller, J.R., Brown, J.D., Lai, C.J., Moon, R.T., 1996. A frizzled homolog functions in a vertebrate Wnt signaling pathway. Curr Biol 6, 1302–1306.

Zaret, K., 1999. Developmental competence of the gut endoderm: genetic potentiation by GATA and HNF3/fork head proteins. Developmental biology 209, 1–10.

