## Supplementary Figure 1 for "Chromatin accessibility and histone acetylation in the regulation of competence in early development"

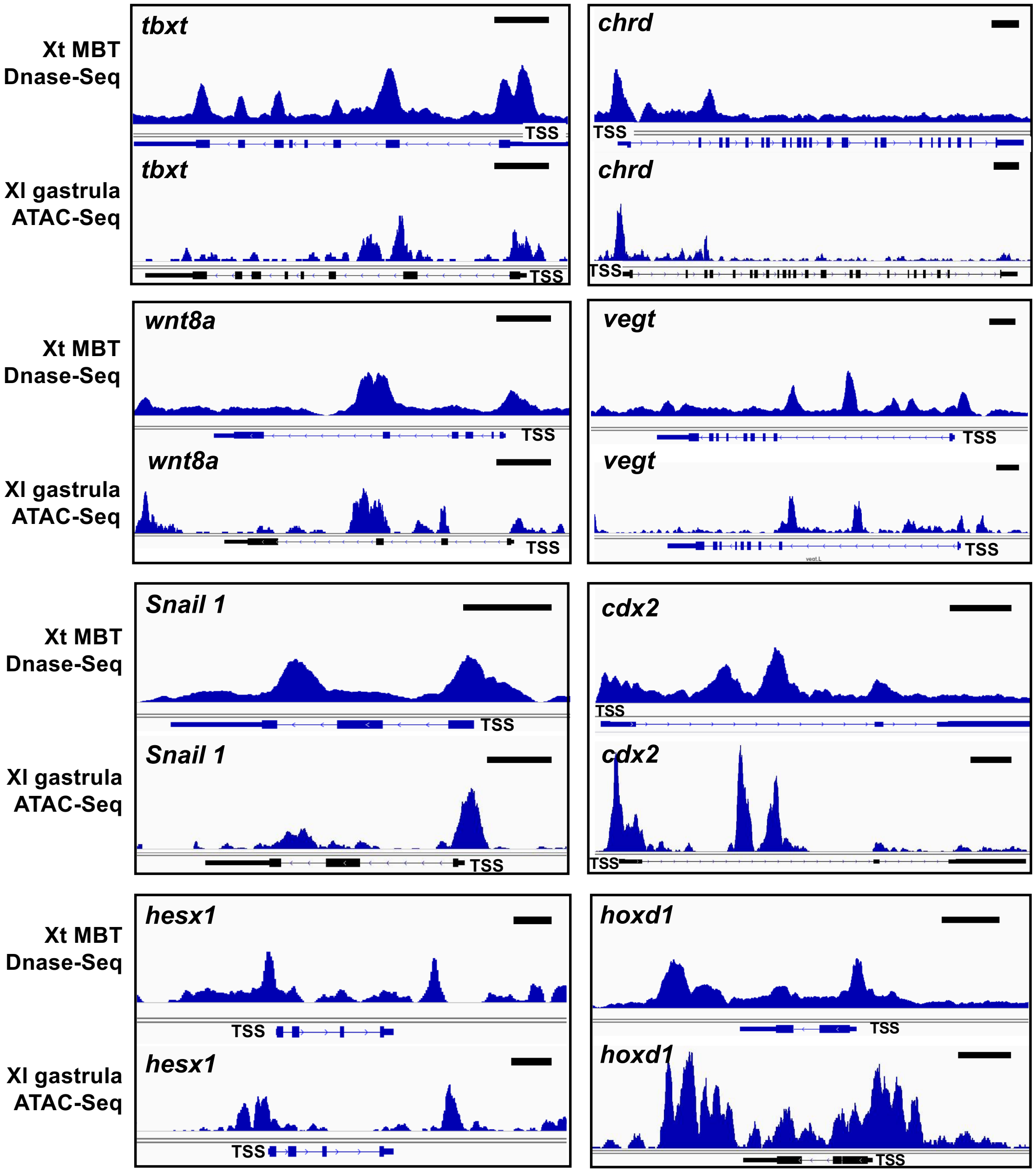


**Chromatin accessibility by ATAC-Seq (*X. laevis* gastrula) vs Dnase-Seq (*X. tropicalis*, MBT)**
ATAC-Seq tracks from stage 10 (early gastrula) *Xenopus laevis* ectodermal explants (GSE GSE76059) were compared to Dnase-Seq tracks from stage 8.5 (MBT) *Xenopus tropicalis* embryos from (Gentsch et al 2019; GSE113186). Tracks were viewed in IGV. Scale bars indicate 1kb. Y-axes were adjusted for each gene to allow side-by-side comparisons and do not reflect relative intensity of signals between different genes, species, stage, or method of analysis. TSS: transcription start site. “Xl”: *Xenopus laevis*; “Xt: *Xenopus topicalis*.
